# Polygenic adaptation: From sweeps to subtle frequency shifts

**DOI:** 10.1101/450759

**Authors:** Ilse Höllinger, Pleuni S Pennings, Joachim Hermisson

**Affiliations:** Mathematics and BioSciences Group, Faculty of Mathematics and Max F. Perutz Laboratories, University of Vienna, Vienna, Austria; Vienna Graduate School of Population Genetics, Vienna, Austria; Department of Biology, San Francisco State University, California, USA

## Abstract

1

Evolutionary theory has produced two conflicting paradigms for the adaptation of a polygenic trait. While population genetics views adaptation as a sequence of selective sweeps at single loci underlying the trait, quantitative genetics posits a collective response, where phenotypic adaptation results from subtle allele frequency shifts at many loci. Yet, a synthesis of these views is largely missing and the population genetic factors that favor each scenario are not well understood. Here, we study the architecture of adaptation of a binary polygenic trait (such as resistance) with negative epistasis among the loci of its basis. The genetic structure of this trait allows for a full range of potential architectures of adaptation, ranging from sweeps to small frequency shifts. By combining computer simulations and a newly devised analytical framework based on Yule branching processes, we gain a detailed understanding of the adaptation dynamics for this trait. Our key analytical result is an expression for the joint distribution of mutant alleles at the end of the adaptive phase. This distribution characterizes the polygenic pattern of adaptation at the underlying genotype when phenotypic adaptation has been accomplished. We find that a single compound parameter, the population-scaled background mutation rate Θ_*bg*_, explains the main differences among these patterns. For a focal locus, Θ_*bg*_ measures the mutation rate at all redundant loci in its genetic background that offer alternative ways for adaptation. For adaptation starting from mutation-selection-drift balance, we observe different patterns in three parameter regions. Adaptation proceeds by sweeps for small Θ_*bg*_ ≾ 0.1, while small polygenic allele frequency shifts require large Θ_bg_ ≿ 100. In the large intermediate regime, we observe a heterogeneous pattern of partial sweeps at several interacting loci.

**Author summary:** It is still an open question how complex traits adapt to new selection pressures. While population genetics champions the search for selective sweeps, quantitative genetics proclaims adaptation via small concerted frequency shifts. To date the empirical evidence of clear sweep signals is more scarce than expected, while subtle shifts remain notoriously hard to detect. In the current study we develop a theoretical framework to predict the expected adaptive architecture of a simple polygenic trait, depending on parameters such as mutation rate, effective population size, size of the trait basis, and the available genetic variability at the onset of selection. For a population in mutation-selection-drift balance we find that adaptation proceeds via complete or partial sweeps for a large set of parameter values. We predict adaptation by small frequency shifts for two main cases. First, for traits with a large mutational target size and high levels of genetic redundancy among loci, and second if the starting frequencies of mutant alleles are more homogeneous than expected in mutation-selection-drift equilibrium, e.g. due to population structure or balancing selection.

## 3 Introduction

Rapid phenotypic adaptation of organisms to all kinds of novel environments is ubiquitous and has been described and studied for decades [1, 2]. However, while the macroscopic changes of phenotypic traits are frequently evident, their genetic and genomic underpinnings are much more difficult to resolve. Two independent research traditions, molecular population genetics and quantitative genetics, have coined two opposite views of the adaptive process on the molecular level: adaptation either by selective sweeps or by subtle allele frequency shifts (sweeps or *shifts* from here on).

On the one hand, population genetics works bottom-up from the dynamics at single loci, without much focus on the phenotype. The implicit assumption of the sweep scenario is that selection on the trait results in sustained directional selection also on the level of single underlying loci. Consequently, we can observe phenotypic adaptation at the genotypic level, where selection drives allele frequencies at one or several loci from low values to high values. Large allele frequency changes are the hallmark of the sweep scenario. If these frequency changes occur in a short time interval, conspicuous diversity patterns in linked genomic regions emerge: the footprints of hard or soft selective sweeps [3–6].

On the other hand, quantitative genetics envisions phenotypic adaptation top-down, from the vantage point of the trait. At the genetic level, it is perceived as a collective phenomenon that cannot easily be broken down to the contribution of single loci. Indeed, adaptation of a highly polygenic trait can result in a myriad of ways through “infinitesimally” small, correlated changes at the interacting loci of its basis (e.g. [1,7,8]). Conceptually, this view rests on the infinitesimal model by Fisher (1918) [9] and its extensions (e.g. [10]). Until a decade ago, the available moderate sample sizes for polymorphism data had strongly limited the statistical detectability of small frequency shifts. Therefore, the detection of sweeps with clear footprints was the major objective for many years. Since recently, however, huge sample sizes (primarily of human data) enable powerful genome-wide association studies (GWAS) to resolve the genomic basis of polygenic traits. Consequently, following conceptual work by Pritchard and coworkers [7, 11], there has been a shift in focus to the detection of polygenic adaptation from subtle genomic signals (e.g. [12–14], reviewed in [15]). Very recently, however, some of the most prominent findings of polygenic adaptation in human height have been challenged [16, 17]. As it turned out, the methods are highly sensitive to confounding effects in GWAS data due to population stratification.

While discussion of the empirical evidence is ongoing, the key objective for theoretical population genetics is to clarify the conditions (mutation rates, selection pressures, genetic architecture) under which each adaptive scenario, sweeps, shifts – or any intermediate type – should be expected in the first place. Yet, the number of models in the literature that allow for a comparison of alternative adaptive scenarios at all is surprisingly limited (see also [18]). Indeed, quantitative genetic studies based on the infinitesimal model or on summaries (moments, cumulants) of the breeding values do not resolve allele frequency changes at individual loci(e.g. [19–22]). In contrast, sweep models with a single locus under selection in the tradition of Maynard Smith and Haigh [3], or models based on adaptive walks or the adaptive dynamics framework (e.g. [23–25]) only allow for adaptive substitutions or sweeps. A notable exception is the pioneering study by Chevin and Hospital [26]. Following Lande [27], these authors model adaptation at a single major quantitative trait locus (QTL) that interacts with an ‘‘infinitesimal background” of minor loci, which evolves with fixed genetic variance. Subsequent models [28, 29] trace the allele frequency change at a single QTL in models with 2-8 loci. Still, these articles do not discuss polygenic adaptation patterns. Most recently, Jain and Stephan [30, 31] studied the adaptive process for a quantitative trait under stabilizing selection with explicit genetic basis. Their analytical approach allows for a detailed view of allele frequency changes at all loci without constraining the genetic variance. However, the model is deterministic and thus ignores the effects of genetic drift. Below, we study a polygenic trait that can adapt via sweeps or shifts under the action of all evolutionary forces in a panmictic population (mutation, selection, recombination and drift). Our model allows for comprehensive analytical treatment, leading to a multi-locus, non-equilibrium extension of Wright’s formula [32] for the joint distribution of allele frequencies at the end of the adaptive phase. This way, we obtain predictions concerning the adaptive architecture of polygenic traits and the population genetic variables that delimit the corresponding modes of adaptation.

The article is organized as follows. The Model section motivates our modeling decisions and describes the simulation method. We also give a brief intuitive account of our analytical approach. In the Results part, we describe our findings for a haploid trait with linkage equilibrium among loci. All our main conclusions in the Discussion part are based on the results displayed here. Further model extensions and complications (diploids, linkage, and alternative starting conditions) are relegated to appendices. Finally, we describe our analytical approach and derive all results in a comprehensive Mathematical Appendix. For the ease of reading, we have tried to keep both the main text and the Mathematical Appendix independent and largely self-contained.

## 4 Model

In the current study, we aim for a “minimal model” of a trait that allows us to clarify which evolutionary forces favor sweeps over shifts and vice versa (as well as any intermediate patterns). For shifts, alleles need to be able to hamper the rise of alleles at other loci via negative epistasis for fitness, e.g. diminishing returns epistasis. Indeed, otherwise one would only observe parallel sweeps. Negative fitness epistasis is frequently found in empirical studies (e.g. [33]) and implicit to the Gaussian selection scheme used by (e.g. [26, 30, 31]). More fundamentally, diminishing returns are a consequence of partial or complete redundancy of genetic effects across loci or gene pathways. Adaptive phenotypes (such as pathogen resistance or a beneficial body coloration) can often be produced in many alternative ways, such that redundancy is a common characteristic of beneficial mutations.

As our basic model, we focus on a haploid population and study adaptation for a polygenic, binary trait with full redundancy of effects at all loci. We assume a non-additive genotype-phenotype map where any single mutation switches the phenotype from its ancestral state (e.g. “non-resistant”) to the adaptive state (“resistant”). Further mutations have no additional effect. On the population level, adaptation can be produced by a single locus where the beneficial allele sweeps to fixation, or by small frequency shifts of alleles at many different loci in different individuals - or any intermediate pattern. The symmetry among loci(no build-in advantage of any particular locus) and complete redundancy of locus effects provides us with a trait architecture that is favorable for collective adaptation via small shifts - and with a modeling framework that allows for analytical treatment. The same model has been used in a preliminary simulation study [6]. In the context of parallel adaptation in a spatially structured population, analogous model assumptions with redundant loci have been used [34–36]. In a second step, we extend our basic model to relax the redundancy condition, as described below.

### 4.1 Basic model

Consider a panmictic population of *N_e_* haploids, with a binary trait *Z* (with phenotypic states *Z*_0_ “non-resistant” and *Z*_1_ “resistant”, see Fig 1). The trait is governed by a polygenic basis of *L* bi-allelic loci with arbitrary linkage (we treat the case of linkage equilibrium in the main text and analyze the effects of linkage in Appendix A.1). Only the genotype with the ancestral alleles at all loci produces phenotype *Z*_0_, all other genotypes produce *Z*_1_, irrespective of the number of mutations they carry. Loci mutate at rate *μ_i_*, 1 ≤ *i* ≤ *L*, per generation (population mutation rate at the *i*th locus: 2*N_e_μ_i_* = Θ_*i*_) from the ancestral to the derived allele. We ignore back mutation. The mutant phenotype *Z*_1_ is deleterious before time *t* = 0, when the population experiences a sudden change in the environment (e.g. arrival of a pathogen). *Z*_1_ is beneficial for time *t* > 0. The Malthusian (logarithmic) fitness function of an individual with phenotype *Z* reads

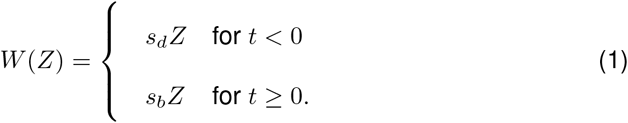

**Fig 1.**
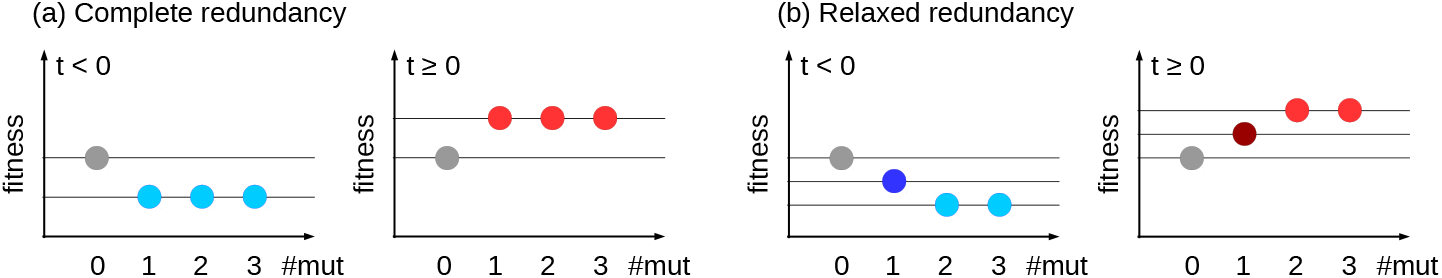
Fitness schemes. The fitness of individuals carrying 0,1,2,3… mutations (y-axis) are given for the complete redundancy (a) and relaxed redundancy (b) model, respectively. Grey balls show the fitness of ancestral wildtype individuals (without mutations). Colored balls represent individuals carrying at least one mutation, for time points *t* < 0 before the environmental change in blue and for *t* ≥ 0 in red.

Without loss of generality, we can assume *Z*_0_ = 0 and *Z*_1_ = 1. We then have *W*(*Z*_0_) = 0. Furthermore, *W*(*Z*_1_) = *s_d_* < 0, respectively *W*(*Z*_1_) = *s_b_* > 0, measure the strength of directional selection on *Z* (e.g. cost and benefit of resistance) before and after the environmental change. For the basic model, we assume that the population is in mutation-selection-drift equilibrium at time *t* = 0.

### 4.2 Model extensions

We extend the basic model in several directions. This includes linkage (Appendix A.1), alternative starting conditions at time *t* = 0 (Appendix A.2), diploids (Appendix A.3), and arbitrary time-dependent selection *s*(*t*) (Mathematical Appendix M.1). Here, we describe how we relax the assumption of complete redundancy of all loci. Diminishing returns epistasis, e.g. due to Michaelis-Menten enzyme kinetics, will frequently not lead to complete adaptation in a single step, but may require multiple steps before the trait optimum is approached. In a model of incomplete redundancy, we thus assume that a first beneficial mutation only leads to partial adaptation. We thus have three states of the trait, the ancestral state for the genotype without mutations, *Z*_0_ = 0 (non-resistant), a phenotype *Z_δ_* = *δ* (partially resistant) for genotypes with a single mutation, and the mutant state *Z*_1_ = 1 (fully resistant) for all genotypes with at least two mutations, see Fig 1(b). For diminishing returns epistasis, we require 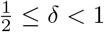. The fitness function is as in Eq (1). A model with asymmetries in the single-locus effects is discussed in Appendix A.4.

### 4.3 Simulation model

For the models described above, we use Wright-Fisher simulations for a haploid, panmictic population of size *N_e_*, assuming linkage equilibrium between all *L* loci in discrete time. Selection and drift are implemented by independent weighted sampling based on the marginal fitnesses of the ancestral and mutant alleles at each locus. Due to linkage equilibrium, the marginal fitnesses only depend on the allele frequencies and not genotypes. Ancestral alleles mutate with probability *μ_i_* per generation at locus i. We start our simulations with a population that is monomorphic for the ancestral allele at all loci. The population evolves for 8*N_e_* generations under mutation and deleterious selection to reach (approximate) mutation-selection-drift equilibrium. Following [6, 37], we condition on adaptation from the ancestral state and discard all runs where the deleterious mutant allele (at any locus) reaches fixation during this time. (We do not show results for cases with very high mutation rates and weak deleterious selection when most runs are discarded). At the time of environmental change, selection switches from negative to positive and simulation runs are continued until a prescribed stopping condition is reached.

We are interested in the genetic architecture of adaptation – the joint distribution of mutant frequencies across all loci – at the end of the rapid adaptive phase.

Following [31], we define this phase as “the time until the phenotypic mean reaches a value close to the new optimum”. Specifically, we stop simulations when the mean fitness 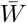 in the population has increased up to a proportion *f_w_* of the maximal attainable increase from the ancestral to the derived state,

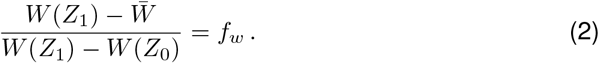

For the basic model with complete redundancy, this simply corresponds to a residual proportion *f_w_* of individuals with ancestral phenotype in the population. Extensions of the simulation scheme to include linkage or diploid individuals are described in Appendices A.1 and A.3.

#### Parameter choices

Unless explicitly stated otherwise, we simulate *N_e_* = 10 000 individuals, with beneficial selection coefficients *s_b_* = 0.1 and 0.01, combined with deleterious selection coefficients *s_d_* = −0.1 and *s_d_* = −0.001 for low and high levels of SGV, respectively. (The corresponding Wrightian fitness values used as sampling weights in discrete time are 1 + *s_b_* and 1 + *s_d_*.) We investigate *L* = 2 to 100 loci. We usually (except in Appendix A.4) assume equal mutation rates at all loci, *μ_i_* = *μ* and define Θ_*l*_ = 2*N_e_μ* as the locus mutation parameter. Mutation rates are chosen such that Θ_*bg*_ := 2*N_e_μ*(*L* − 1) (the background mutation rate, formally defined below in Eq (10)) takes values from 0.01 to 100. We typically simulate 10 000 replicates per mutation rate and stop simulations when the population has reached the new fitness optimum up to *f_w_* = 0.05. In the model with complete redundancy, we thus stop simulations when the frequency of individuals with mutant phenotype *Z*_1_ has increased to 95%. Different stopping conditions are explored in Appendix A.7.

### 4.4 Analytical analysis

We partition the adaptive process into two phases (see Fig 2 for illustration). An initial *stochastic phase*, governed by selection, drift, and mutation describes the origin and establishment of mutant alleles at all loci. We call mutants ‘established” if they are not lost again due to genetic drift. The subsequent *deterministic phase* governs the further evolution of established alleles until the stopping condition is reached as described above. While mutation and drift can be ignored during the deterministic phase, interaction effects due to epistasis and linkage become important (in our model, they enter, in particular, through the stopping condition). We give a brief overview of our analytical approach below. A detailed account with the derivation of all results is provided in the Mathematical Appendix.

**Fig 2.**
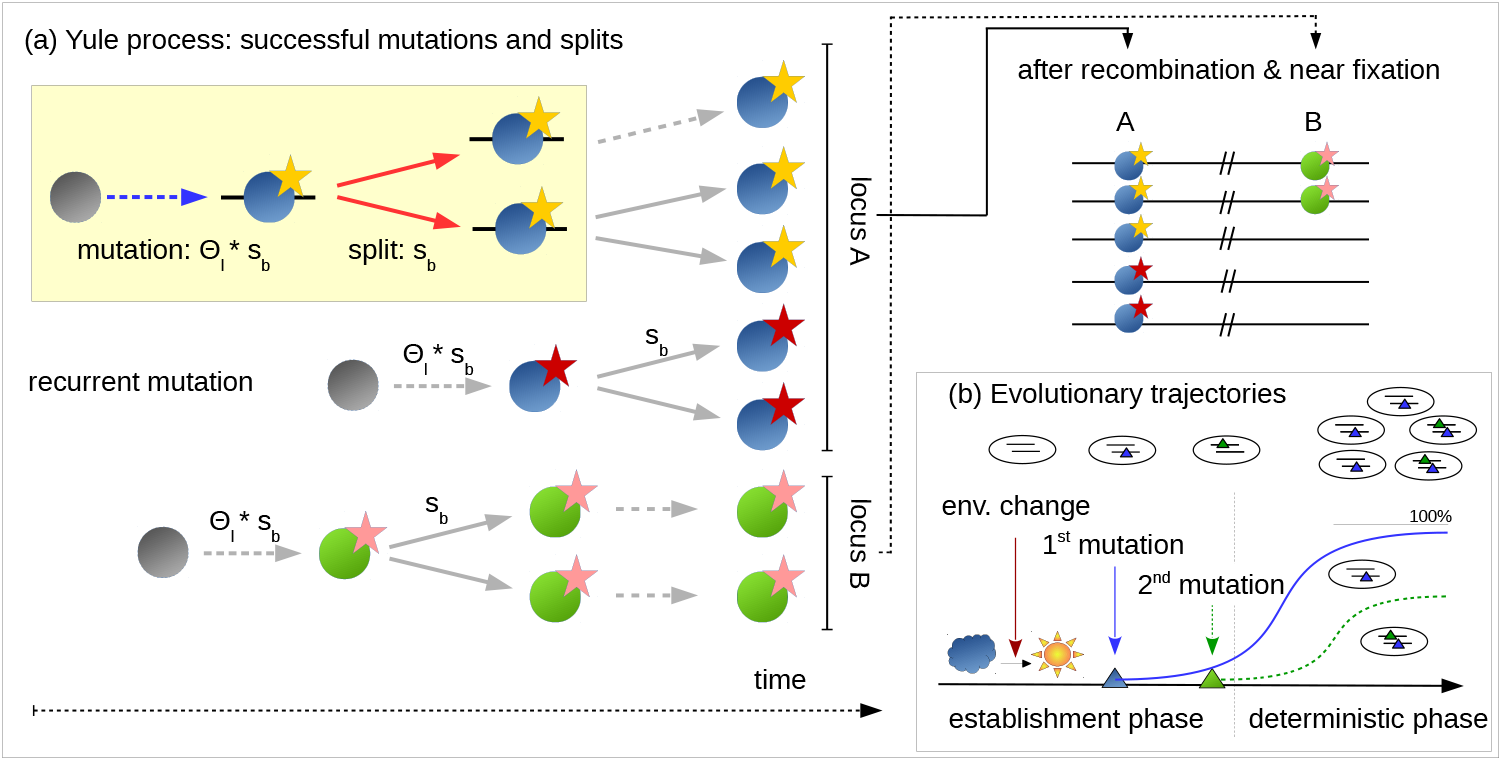
Phases of polygenic adaptation. The adaptive process is partitioned into two phases. The initial, stochastic phase describes the establishment of mutant alleles. Ignoring epistasis during this phase, it can be described by a *Yule* process (panel a), with two types of events (yellow box). Either a new mutation occurs and establishes with rate Θ_*l*_ · *s_b_* or an existing mutant line splits into two daughter lines at rate *s_b_*. Mutations and splits can occur in parallel at all loci of the polygenic basis, (here 2 loci, shown in green and blue). Yellow and red stars at the blue locus indicate establishment of two redundant mutations at this locus. When mutants have grown to higher frequencies, the adaptive process enters its second, deterministic phase, where drift can be ignored (panel b). During the deterministic phase, the trajectories of mutations at different loci constrain each other due to epistasis. We refer to the locus ending up at the highest frequency as the *major* locus (here in blue) and to all others as *minor* loci (here one in green).

During the *stochastic phase*, we model the origin and spread of mutant copies as a so-called *Yule pure birth process* following [38] and [39]. The idea of this approach is that we only need to keep track of mutations that found “immortal lineages”, i.e. derived alleles that still have surviving offspring at the time of observation (see Fig 2 for the case of *L* = 2 loci). Forward in time, new immortal lineages can be created by two types of events: new mutations at all loci start new lineages, while birth events lead to splits of existing lineages into two immortal lineages. For *t* > 0 (after the environmental change), in particular, new mutations at the *i*th locus arise at rate *N_e_μ_i_* per generation and are destined to become established in the population with probability ≈ 2*s_b_*.

Similarly, birth of new immortal lineages due to split events in the Yule process occur at rate *s_b_* (because the selection coefficient measures the excess of births over deaths in the underlying population). For the origin of new immortal lineages in the Yule process and their subsequent splitting we thus obtain the rates

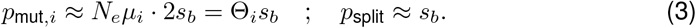

Extended results including standing genetic variation and time-dependent fitness are given in the Appendix. Assume now that there are currently {*k*_1_,… *k_L_*}, 0 ≤ *k_j_* ≪ *N_e_* mutant lineages at the *L* loci. The probability that the next event (which can be a split or a mutation) occurs at locus *i* is

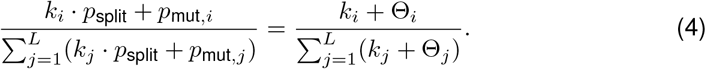

Importantly, all these transition probabilities among states of the Yule process are constant in time and independent of the mutant fitness *s_b_*, which cancels in the ratio of the rates. As the number of lineages at all loci increases, their joint distribution (across replicate realizations of the Yule process) approaches a limit. In particular, as shown in the Appendix, the joint distribution of frequency ratios *x_i_* := *k_i_*/*k*_1_ in the limit *k*_1_ → ∞ is given by an *inverted Dirichlet distribution*

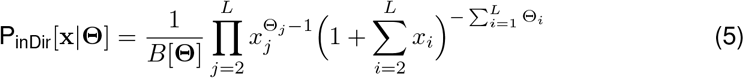

where x = (*x*_2_,…,*x_L_*) and **Θ** = (Θ_1_,…,Θ_*L*_) are vectors of frequency ratios and locus mutation rates, respectively, and where 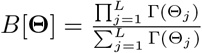 is the generalized Beta function and Γ(*z*) is the Gamma function. Note that Eq (5) depends only on the locus mutation rates, but not on selection strength.

After the initial stochastic phase, the dynamics of established mutant lineages that have evaded stochastic loss can be adequately described by deterministic selection equations. For allele frequencies *p_i_* at locus *i*, assuming linkage equilibrium, we obtain (consult the Mathematical Appendix M.1, Eq (M.2a), for detailed derivations)

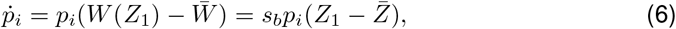

where 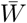 and 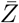 are population mean fitness and mean trait value. For the mutant frequency ratios *x_i_* = *p_i_*/*p*_1_, we obtain

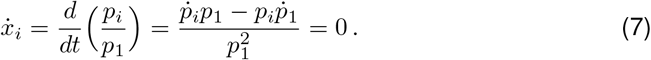

We thus conclude that the frequency ratios *x_i_* do not change during the deterministic phase. In particular, this means that Eq (5) still holds at our time of observation at the end of the rapid adaptive phase. This is even true with linked loci. Finally, derivation of the joint distribution of mutant frequencies pi (instead of frequency ratios *x_i_*) at the time of observation requires a transformation of the density. In general, this transformation depends on the stopping condition *f_w_* and on other factors such as linkage. Assuming linkage equilibrium among all selected loci, we obtain (see the Mathematical Appendix, Theorem 2, Eq (M.20))

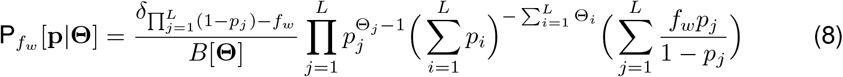

for **p** = (*p*_1_, …,*p_L_*) in the *L*-dimensional hypercube of allele frequencies. The delta function *δ_X_* restricts the distribution to the *L* − 1 dimensional manifold defined via the stopping condition 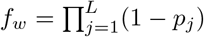. Further expressions, also including linkage, are given in the Mathematical Appendix and in Appendix A.1. In general, the joint distribution corresponds to a family of generalized Dirichlet distributions.

We assess the adaptive architecture not as a function of time, but as a function of progress in phenotypic adaptation, measured by *f_w_*, Eq (2). Hence, *f_w_* rather than time *t* plays the role of a dynamical variable in the joint distribution, see Eq (8). In the special case *f_w_* → 0 (i.e. complete adaptation, enforcing fixation at at least one locus), this distribution is restricted to a boundary face of the allele frequency hypercube and Eq (8) reduces to the inverted Dirichlet distribution given above in Eq (5). In the Results section below, we assess our analytical approximations for the joint distributions of adaptive alleles, Eq (5) and Eq (8), and discuss their implications in the context of scenarios of polygenic adaptation, ranging from sweeps to small frequency shifts.

**Table 1.**
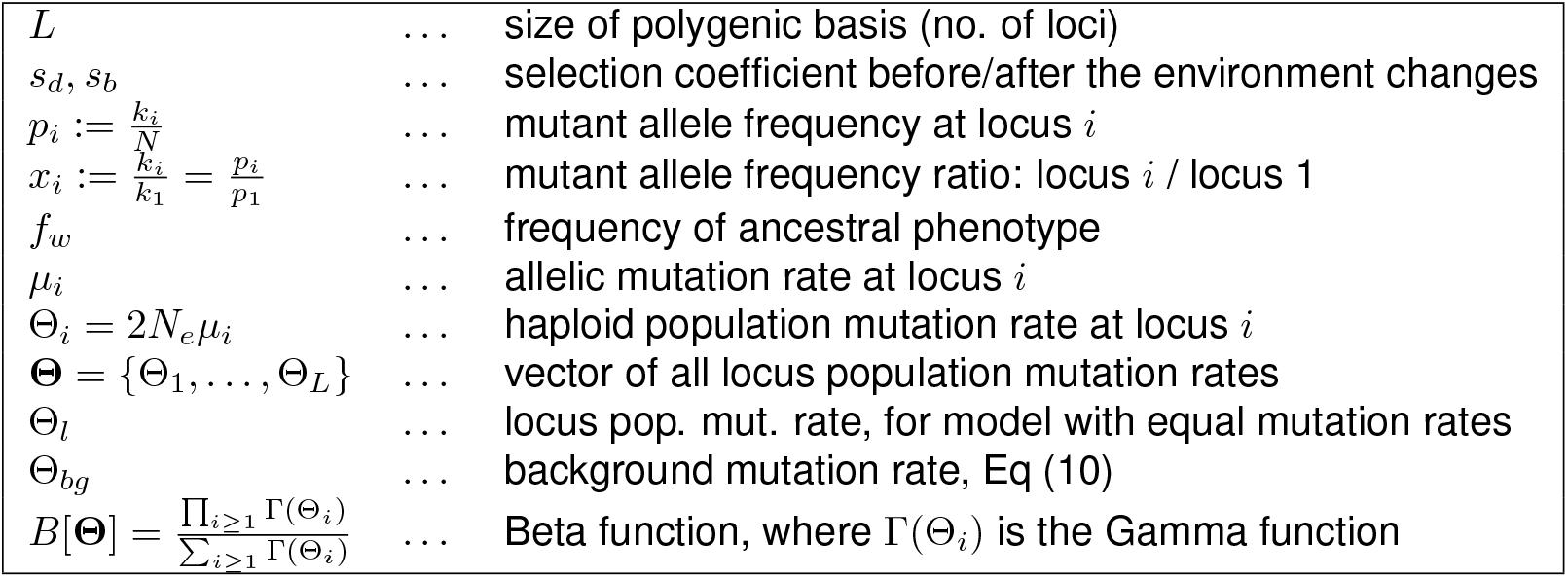
Glossary

|  |  |  |
| --- | --- | --- |
| $L$ | ... | size of polygenic basis (no. of loci) |
| $s_d, s_b$ | ... | selection coefficient before/after the environment changes |
| $p_i := \frac{k_i}{N}$ | ... | mutant allele frequency at locus $i$ |
| $x_i := \frac{k_i}{k_1} = \frac{p_i}{p_1}$ | ... | mutant allele frequency ratio: locus $i$ / locus 1 |
| $f_w$ | ... | frequency of ancestral phenotype |
| $\mu_i$ | ... | allelic mutation rate at locus $i$ |
| $\Theta_i = 2N_e\mu_i$ | ... | haploid population mutation rate at locus $i$ |
| $\Theta = \{\Theta_1, \dots, \Theta_L\}$ | ... | vector of all locus population mutation rates |
| $\Theta_l$ | ... | locus pop. mut. rate, for model with equal mutation rates |
| $\Theta_{bg}$ | ... | background mutation rate, Eq (10) |
| $B[\Theta] = \frac{\prod_{i \geq 1} \Gamma(\Theta_i)}{\sum_{i \geq 1} \Gamma(\Theta_i)}$ | ... | Beta function, where $\Gamma(\Theta_i)$ is the Gamma function |

## 5 Results

While the joint distribution of allele frequencies, Eq (8), provides comprehensive information of the adaptive architecture, low-dimensional summary statistics of this distribution are needed to describe and classify distinct types of polygenic adaptation. To this end, we order loci according to their contribution to the adaptive response. In particular, we call the locus with the highest allele frequency at the stopping condition the *major locus* and all other loci *minor loci*. Minor loci are further ordered according to their frequency (first minor, second minor, etc.). The marginal distributions of the major locus or *k*th minor locus are 1-dimensional summaries of the joint distribution. Importantly, these summaries are still *collective* because the role of any specific locus (its order) is defined through the allele frequencies at *all* loci. This is different for the marginal distribution at a fixed focal locus, which is chosen irrespective of its role in the adaptive process, e.g. [26, 28, 29].

Concerning our nomenclature, note that the *major* and *minor* loci do not differ in their effect size, as they are completely redundant. Still, the major locus is the one with the largest contribution to the adaptive response and would yield the strongest association in a GWAS case-control study.

In the following, we analyze adaptive trait architectures in three steps. In Section 5.1 we use the expected allele frequency ratio of minor and major loci as a one-dimensional summary statistic. Subsequently, in Section 5.2, we analyze the marginal distributions of major and minor loci for a trait with 2 to 100 loci. Finally, in Section 5.3 we investigate the robustness of our results under conditions of relaxed redundancy. Further results devoted to diploids, linkage, asymmetric loci, and alternative starting conditions are provided in the Appendices.

### 5.1 Expected allele frequency ratio

For our biological question concerning the type of polygenic adaptation, the ratio of allele frequency changes of minor over major loci is particularly useful. With “sweeps at few loci”, we expect large differences among loci, resulting in ratios that deviate strongly from 1. In contrast, with “subtle shifts at many loci”, multiple loci contribute similarly to the adaptive response and ratios should range close to 1. Our theory (explained above) predicts that these ratios are the outcome of the stochastic phase, and their distribution is preserved during the deterministic phase. They are thus independent of the precise time of observation. For our results in this section, we assume that the mutation rate at all *L* loci is equal, Θ_*i*_ ≡ Θ_*l*_, for all 1 ≤ *i* ≤ *L*. This corresponds to the symmetric case that is most favorable for a “small shift” scenario. Results for asymmtric mutation rates are reported in Appendix A.4.

Consider first the case of *L* = 2 loci. There is then a single allele frequency ratio “minor over major locus”, which we denote by *x*. For two loci, the joint distribution of frequency ratios from Eq (5) reduces to a *beta-prime* distribution. Conditioning on the case that the first locus is the major locus (probability 1/2 for the symmetric model), we obtain for 0 ≤ *x* ≤ 1,

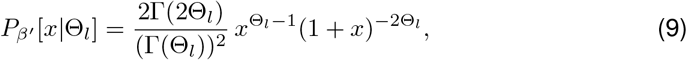

Fig 3 compares the expectation of this analytical prediction with simulation results for a range of parameters for the strength of beneficial selection *s_b_* and for the level of standing genetic variation (SGV implicitly given by the strength of deleterious selection *s_d_* before the environmental change). There are two main observations. First, the simulation results demonstrate the importance of the scaled mutation rate Θ_*bg*_ ≡ Θ_*l*_ (for two loci). Low Θ_*bg*_ leads to sweep-like adaptation (heterogeneous adaptation response among loci, E[*x*] << 1), whereas high Θ_*bg*_ leads to shift-like adaptation (homogeneous response, E[*x*] near 1). Second, the panels show that the selection intensity given by sd and s_b_ has virtually no effect. Both results are predicted by the analytical theory (Eq (9)). In Appendix A.1, we further show that these results hold for arbitrary degrees of linkage (including complete linkage), see Fig S.1.

**Fig 3.**
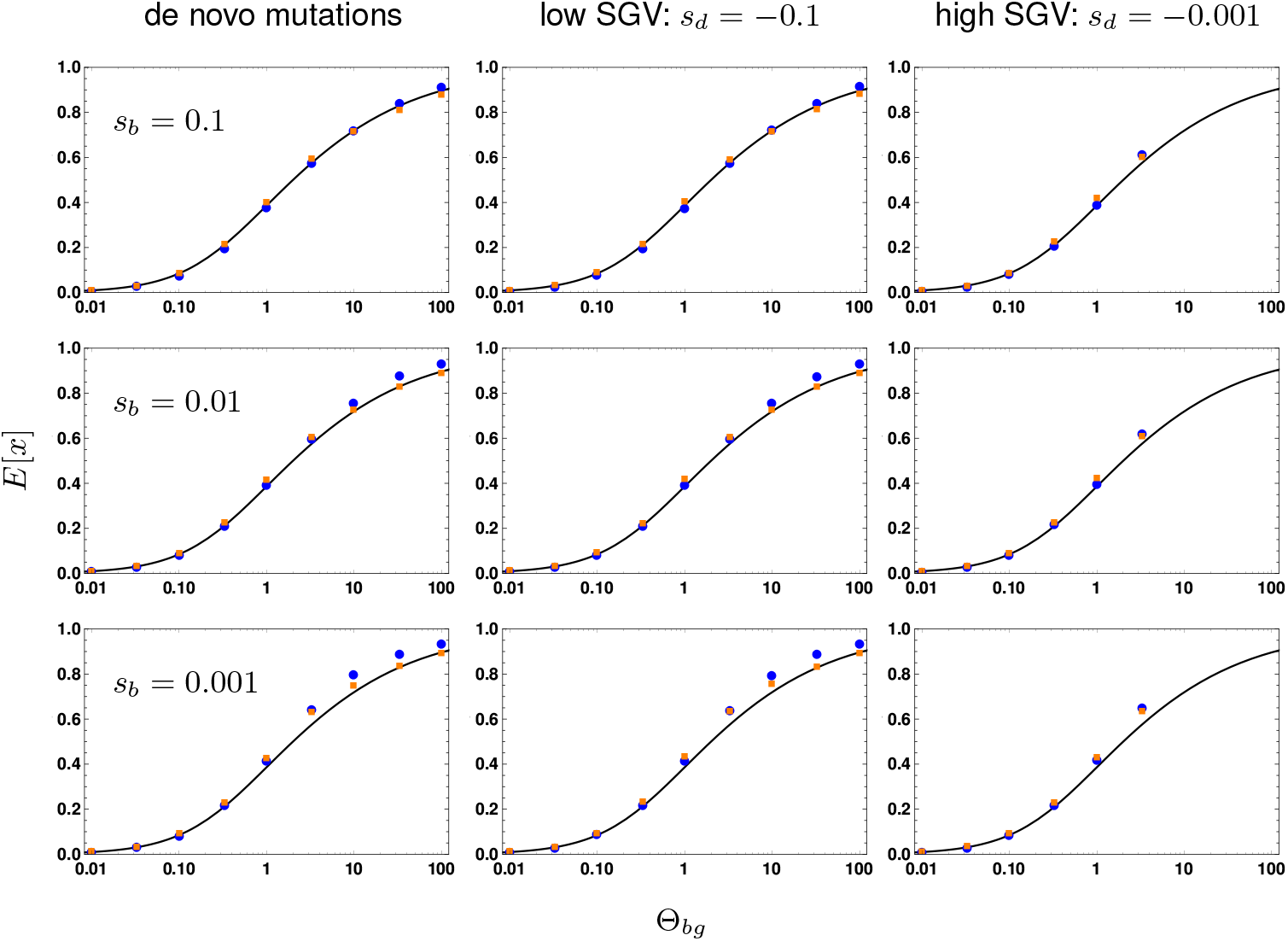
Effect of selection strength and SGV on the frequency ratio E[*x*]. We contrast the expected allele frequency ratios of the first minor locus (with the second highest frequency) over the major locus (with the highest frequency) for 2 loci (blue dots) and for 10 loci (orange dots) with analytical predictions (Appendix, Eq M.16, black curve). E[*x*] is shown as a function of Θ_*bg*_ (= Θ_*l*_ for the 2-locus case). Panels correspond to different strengths of positive selection (*s_b_*, rows) and levels of SGV (no SGV, strongly deleterious *s_d_* = −0.1, weakly deleterious *s_d_* = −0.001, columns). We find that neither factor alters the expected ratio. We do not obtain results for Θ_*bg*_ > 10 and *s_d_* = −0.001, where strong recurrent mutation overwhelmes weak selection, such that mutant alleles fix even before the environmental change. Results for 10 000 replicates, standard errors < 0.005 (smaller than symbols).

For more than two loci, *L* > 2, one-dimensional marginal distributions of the joint distribution, Eq (5), generally require (*L* − 1)-fold integration, which can be complicated. However, it turns out that the key phenomena to characterize the adaptive architecture can still be captured by the 2-locus formalism, with appropriate rescaling of the mutation rate. For the general *L*-locus model, we broaden our definition of the summary statistic x above to describe the allele frequency ratio of the *first minor* locus and the major locus. To relate the distribution of *x* in the *L*-locus model to the one in the 2-locus model, we reason as follows: For small locus mutation rates Θ_*l*_, the order of the loci is largely determined by the order at which mutations that are destined for establishment originate at these loci. *I.e*., the locus where the first mutation originates ends up as the major locus and the first minor locus is usually the second locus where a mutation destined for establishment originates. The distribution of the allele frequency ratio *x* is primarily determined by the distribution of the waiting time for this second mutation after origin of the first mutation at the major locus. In the 2-locus model, this time will be exponentially distributed, with parameter 1/Θ_*l*_. In the *L*-locus model, however, where *L* − 1 loci with total mutation rate Θ_*l*_(*L* − 1) compete for being the “first minor”, the parameter for the waiting-time distribution reduces to 1/(Θ_*l*_(*L* − 1)). We thus see from this argument that the decisive parameter is the cumulative *background mutation rate*

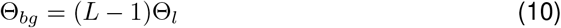

at all minor loci in the background of the major locus. In Fig 3 (orange dots) we show simulations of a *L* = 10 locus model with an appropriately rescaled locus mutation rate Θ_*l*_ → Θ_*l*_/9, such that the background rate Θ_*bg*_ is the same as for the 2-locus model. We see that the analytical prediction based on the 2-locus model provides a good fit for the 10-locus model. A more detailed discussion of this type of approximation is given in Appendix A.5.

### 5.2 Genomic architecture of polygenic adaptation

While the distribution of allele frequency ratios, Eqs (5) and (9), offers a coarse (but robust) descriptor of the adaptive scenario, the joint distribution of allele frequencies at the end of the adaptive phase, Eq (8), allows for a more refined view. In contrast to the distribution of ratios, the results now depend explicitly on the stopping condition (the time of observation) and on linkage among loci. We assume linkage equilibrium in this section and assess the mutant allele frequencies when the frequency of the remaining wildtype individuals in the population is *f_w_* (= 0.05 in our figures) has dropped to a fixed value of *f_w_* = 0.05. In Appendix A.7, we complement these results and study the changes in the adaptive architecture when *f_w_* is varied.

Fig 4 displays the main result of this section. It shows the marginal distributions of all loci, ordered according to their allele frequency at the time of observation (major locus, 1st, 2nd, 3rd minor locus, *etc*.) for traits with *L* = 2, 10, 50, and 100 loci. Panels in the same row correspond to equal background mutation rates Θ_*bg*_ = (*L* − 1)Θ_*l*_, but note that the locus mutation rates Θ_*l*_ are not equal. The figure reveals a striking level of uniformity of adaptive architectures with the same Θ_*bg*_, but vastly different number of loci. For Θ_*bg*_ ≤ 1 (the first three rows), the marginal distributions for lociof the same order (same color in the Figure) across traits with different *L* is almost invariant. For large Θ_*bg*_, they converge for sufficiently large *L* (e.g. for Θ_*bg*_ = 10, going from *L* = 10 to *L* = 50 and to L = 100). In particular, the background mutation rate Θ_*bg*_ determines the shape of the major-locus distribution (red in the Figure) for high *p* → 1 − *f_w_* = 0.95 (the maximum possible frequency, given the stopping condition). For Θ_*bg*_ < 1, this distribution is sharply peaked with a singularity at *p* = 1 − *f_w_*, whereas it drops to zero for high *p* if Θ_*bg*_ > 1 (see also the analytical results below).

**Fig 4.**
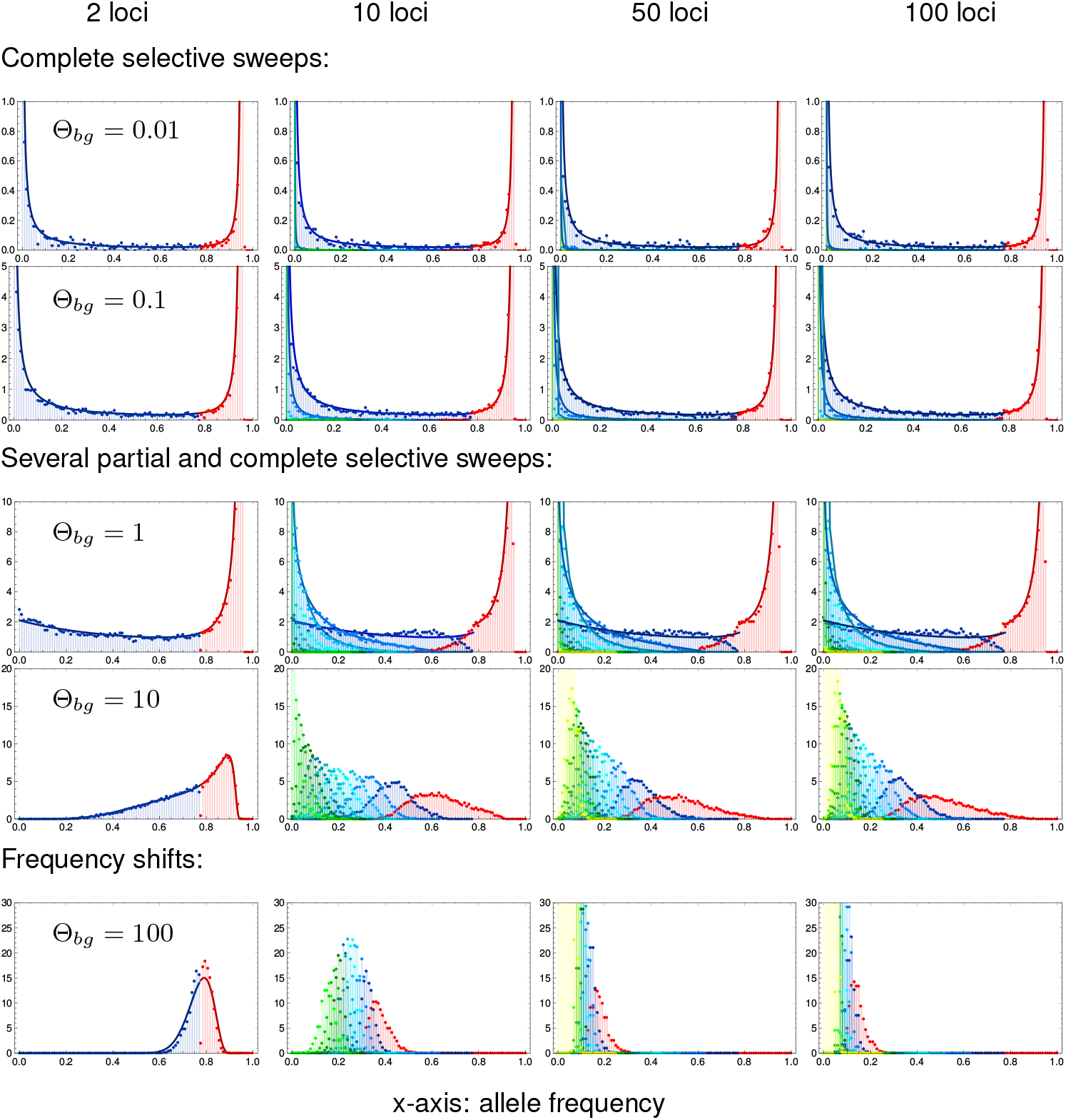
Genomic architecture of polygenic adaptation. We distinguish three patterns of architectures with increasing genomic background mutation rate θ_*bg*_: complete sweeps, for θ_*bg*_ ≾ 0.1, heterogeneous partial sweeps at several loci for 0.1 < θ_*bg*_ < 100, and polygenic frequency shifts for θ_*bg*_ > 100. The plots show the marginal distributions of all loci, ordered according to their allele frequency, i.e. the major locus in red and all following (first, second, third, etc. minors) in blue to green to yellow. Lines in respective colors show analytical predictions, Appendix A.5. Simulations were stopped once the populations have adapted to 95% of the maximum mean fitness in each of 10000 replicates, resulting in an the upper bound for the major locus distribution at, *p*_1_ = 0.95. Simulations for *s_b_* = −*s_d_* = 0.1. Note the different scaling of the y-axis (density, normalized to 1 per locus) for different mutation rates.

As predicted by the theory, Eq (8) and below, simulations confirm that the overall selection strength does not affect the adaptive architecture (see supplementary Fig S.11 for comparison of simulation results for *s_b_* = 0.1 and *s_b_* = 0.01). As discussed in Appendix A.1, sufficiently tight linkage does change the shape of the distributions. Importantly, however, it does not affect the role of Θ_*bg*_ in determining the singularity of the major-locus distribution. This confirms the key role of the background mutation rate as a single parameter to determine the adaptive scenario in our model. While Θ_*bg*_ = 1 separates architectures that are dominated by a single major locus (Θ_*bg*_ < 1) from collective scenarios (with Θ_*bg*_ > 1), the classical sweep or shift scenarios are only obtained if Θ_*bg*_ deviates strongly from 1. We therefore distinguish three adaptive scenarios.

- Θ_*bg*_ ≾ 0.1, *single completed sweeps*. For Θ_*bg*_ ≪ 1 (first two rows of Fig 4), the distribution of the major locus is concentrated at the maximum of its range, while all other distributions are concentrated around 0. Adaptation thus occurs at a single locus, via a selective sweep from low to high mutant frequency. Contributions by further loci are rare. If they occur at all they are usually due to a single runner-up locus (the highest minor locus).
- 0.1 < Θ_*bg*_ < 100, *heterogeneous partial sweeps*. With intermediate background mutation rates (third and forth row of Fig 4), we still observe a strong asymmetry in the frequency spectrum. Even for Θ_*bg*_ = 10, there is a clear major locus discernible, with most of its distribution for *p* > 0.5. However, there is also a significant contribution of several minor loci that rise to intermediate frequencies. We thus obtain a heterogeneous pattern of partial sweeps at a limited number of loci.
- Θ_*bg*_ ≿ 100, *homogeneous frequency shifts*. Only for high background mutations rates Θ_*bg*_ > 1 (last row of Fig 4 with Θ_*bg*_ = 100), the heterogeneity in the locus contributions to the adaptive response vanishes. There is then no dominating major locus. For only 2 loci, these shifts are necessarily still quite large, but for traits with a large genetic basis (large *L*; the only realistic case for high values of Θ_*bg*_), adaptation occurs via subtle frequency shifts at many loci.

#### Analytical predictions

To gain deeper understanding of the polygenic architecture – and for quantitative predictions – we dissect our analytical result for the joint frequency spectrum in Eq (8). We start with the case of *L* = 2 loci, allowing for different locus mutation rates Θ_1_ and Θ_2_. The marginal distribution at the first locus reads (from Eq (8), after integration over *p*_2_),

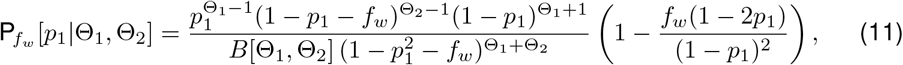

for 0 ≤ *p*_1_ ≤ 1 − *f_w_* (see also Appendix A.6). The distribution has a singularity at *p*_1_ = 0 if the corresponding *locus* mutation rate is smaller than one, Θ_1_ < 1. It has a singularity at *p*_1_ = 1 − *f_w_* if the corresponding *background* mutation rate (which is just the mutation rate at the other locus for *L* = 2) is smaller than one, Θ_2_ < 1. The marginal distributions at the major locus, 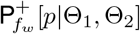, and the minor locus, 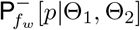, follow from Eq (11) as

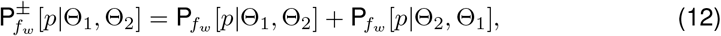

where 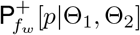 is defined for 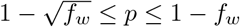 and 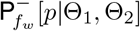 is defined for 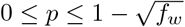. The sum in Eq (12) accounts for the alternative events that either the first or the second locus may end up as the major (or minor) locus. Consequently, 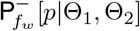 has a singularity at *p* = 0 if the *minimal locus mutation rate* Θ_*l*_ = min[Θ_1_, Θ_2_] < 1. Analogously, 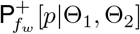 has a singularity at *p* = 1 − *f_w_* if the *minimal background mutation rate* Θ_*bg*_ = min[Θ_1_, Θ_2_] < 1. The left column of Fig 4 shows the distributions at the major and minor locus for *L* = 2 in the symmetric case Θ_1_ = Θ_2_ = Θ_*l*_ = Θ_*bg*_ and *f_w_* = 0.05. Simulations for a population of size *N_e_* = 10 000 and analytical predictions match well.

How do these results generalize for *L* > 2? We again allow for unequal locus mutation rates Θ_*i*_. It is easy to see from Eq (8) that the marginal distribution at the ith locus has a singularity at *p_i_* = 0 for Θ_*i*_ < 1. In the Mathematical Appendix M.3, we further show that it has a second singularity at *p_i_* = 1 − *f_w_* if the corresponding background mutation rate 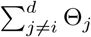 is smaller than 1. As a first step, we split the joint distribution, Eq (8), into the marginal distribution at the major locus 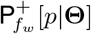 (defined for 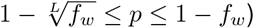) and a cumulative distribution at all other (minor) loci, 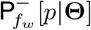 (defined for 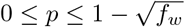). Since any locus can end up as the major locus (with probability > 0), 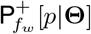 has a singularity at *p* = 1 − *f_w_* for

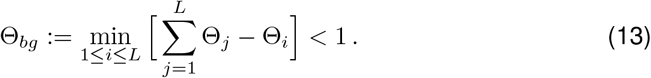

This equation generalizes the definition of the background mutation rate, Eq (10), to the case of unequal locus mutation rates. Similarly, 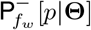 has a singularity at *p* = 0 if

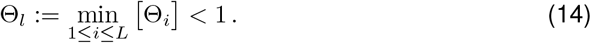

As long as Θ_*bg*_ ≤ 1, we can approximate both the major-locus distribution 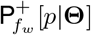 and the cumulative minor locus distribution 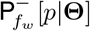 for arbitrary *L* by formulas for a 2-locus model with locus mutation rates matching Θ_*l*_ and Θ_*bg*_ of the multi-locus model, Eq (12). Similarly, we can use results from a *k*-locus model to match the marginal distributions of the largest *k* loci (i.e., up to the (*k* − 1)th minor) in models with *L* > *k* loci, upon rescaling of the mutation rates. As explained for the ratio of the first minor and major locus in the previous section, rescaling rules match the expected waiting time for a mutation (destined for establishment) at the *k*th locus after the origin of a first mutation. Details are given in the Appendix A.5. In Fig 4, we use formulas derived from a *k*-locus model (*k* ≤ 4) to approximate the (*k* − 1)st minor locus distribution of models with *L* = 10; 50; 100 lociand Θ_*bg*_ ≤ 1. These approximations work well as long as these leading loci dominate the adaptive architecture of the trait, which is the case for Θ_*bg*_ ≤ 1.

### 5.3 Relaxing complete redundancy

To complete our picture of adaptive architectures, we investigate the robustness of our model assumption against relaxation of redundancy. As explained above (*Model extensions* and Fig 1), we implement diminishing returns epistasis, such that an individual with a single mutation has fitness δ*s_b_*/_*d*_, while individuals carrying more than one mutation have fitness *s*_*b*/*d*_. With small deviations from complete redundancy (e.g. *δ* = 0.9, stopping at 5% ancestral phenotypes, see Fig S.10) we obtain basically no differences in the genomic patterns of adaptation. With larger deviations (e.g. *δ* = 0.5) quantitative differences appear. However, the qualitative picture concerning the scenario of polygenic adaptation remains the same.

Fig 5 shows the marginal frequency distributions of major and minor loci for a trait with relaxed redundancy with *δ* = 0.5 that is sampled when the population has accomplished 95% of the fitness increase on its way to the new optimum, Eq (2). Given the fitness function, this is not possible with adaptation at only a single locus. At least two loci are needed. The Figure compares the simulation data for the relaxed redundancy model (colored dots) and the full redundancy model (dots in back and gray). As in Fig 4, traits in the same row have the same background mutation rate Θ_*bg*_. However, the background rate for the model with relaxed redundancy is redefined as

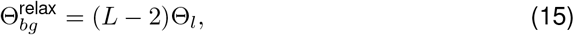

where Θ_*l*_ is the locus mutation rate (equal at all loci). We thus define the background rate, more precisely, as the combined population-scaled mutation rate of all loci *that are not essential* to accomplish adaptation of the phenotype and, thus, are truly redundant. With this choice, the adaptive architecture of the relaxed redundancy model reproduces the one of the model with full redundancy – up to a shift in the number of the loci due to an extra locus that is needed for adaptation with relaxed redundancy. The Figure captures this by comparing traits with relaxed redundancy with *L* = 3,4,11, and 101 loci to fully redundant traits with one fewer locus. The inset figures in the column for *L* = 4 loci show the same scenario, but with an *averaged* marginal distribution for the two largest loci with relaxed redundancy (in green).

**Fig 5.**
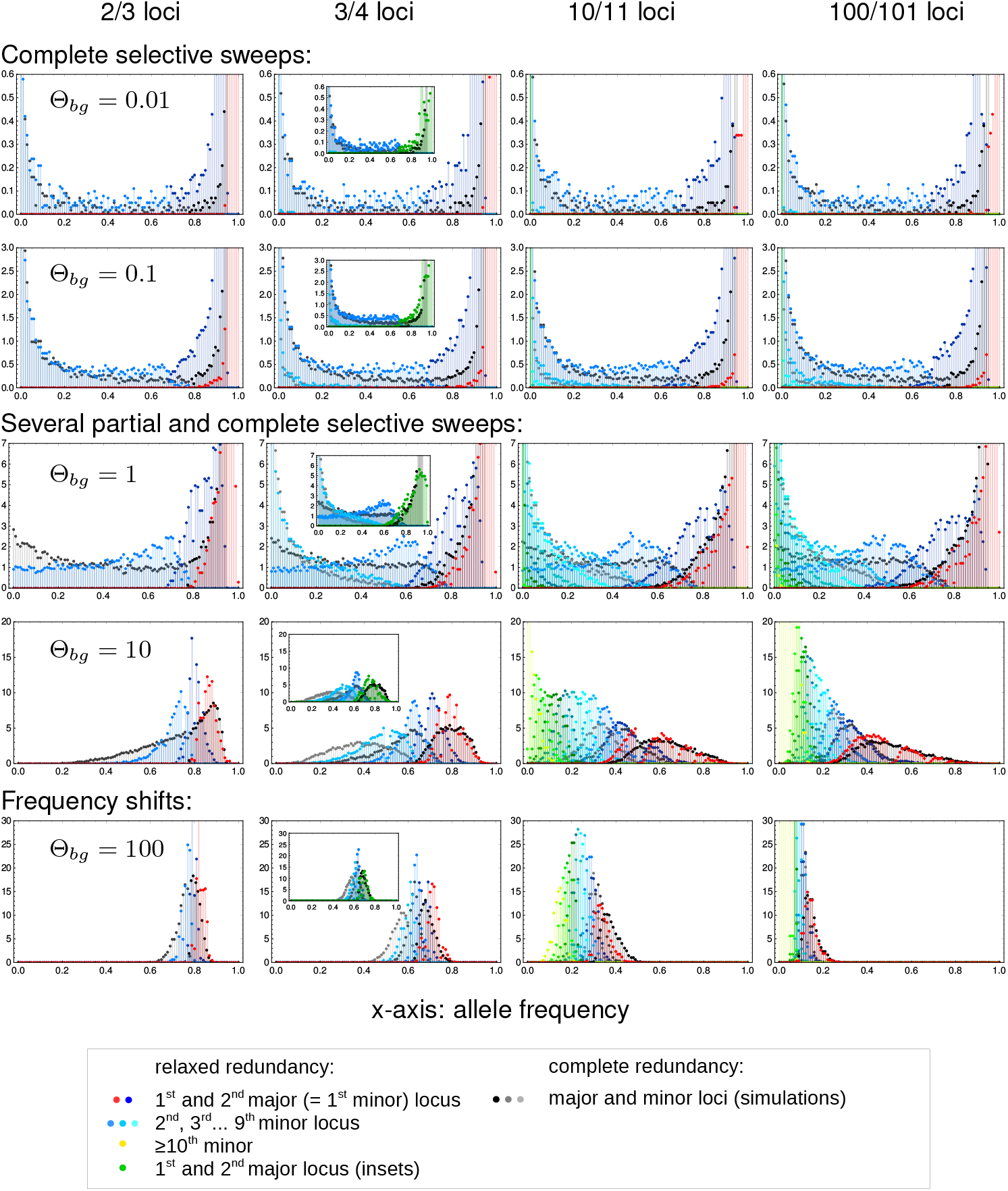
Relaxed redundancy. Relaxing redundancy such that a single mutant has fitness 1 + 0.5*s*_*b*/*d*_ and only two mutations or more confer the full fitness effect (1 + *s*_*b*/*d*_) demonstrates the robustness of our model. As in Fig 4, allele frequency distributions of derived alleles are displayed once the population has reached 95% of maximum attainable mean population fitness. Genomic patterns of adaptation show similar characteristics as with complete redundancy. Due to relaxed redundancy, an additional ‘‘major locus” is required to reach the adaptive optimum. As explained in the main text, the distribution of the *k*th largest locus with complete redundancy therefore corresponds to the distribution of the *k* + 1st largest locus with relaxed redundancy. Insets in the second column show the same data with the distributions of the two major loci for relaxed redundancy combined (in green).

- For mutation rates, Θ_*bg*_ ≪ 1, we still find adaptation by sweeps. Relative to the full redundancy model, we now observe two “major” sweep loci instead of only a single sweep. The inset (for *L* = 4) shows that their averaged distributions matches the major locus distribution of the full redundancy model. The distribution at the third largest locus (the “first minor” locus with relaxed redundancy) resembles the corresponding distribution of the first minor locus of the trait with full redundancy.
- For intermediate mutation rates, 0.1 < Θ_*bg*_ < 100, the pattern is dominated by partial sweeps. We clearly see the similarity in the marginal distributions of the kth largest locus with full redundancy and the *k* + 1st largest locus of the relaxed redundancy trait. For the two major loci with relaxed redundancy, we again see (inset) that the averaged distribution matches the major-locus distribution of the full redundancy model.
- Finally, for strong mutation, Θ_*bg*_ ≿ 100, adaptation again occurs by small frequency shifts at many loci.

In summary, our results show that relaxing redundancy leads to qualitatively similar results, but with a reduced “effective” background mutation rate that only accounts for “truly redundant” loci.

## 6 Discussion

Traits with a polygenic basis can adapt in different ways. Few or many loci can contribute to the adaptive response. The changes in the allele frequencies at these loci can be large or small. They can be homogeneous or heterogeneous. While molecular population genetics posits large frequency changes – selective sweeps – at few loci, quantitative genetics views polygenic adaptation as a collective response, with small, homogeneous allele frequency shifts at many loci. Here, we have explored the conditions under which each adaptive scenario should be expected, analyzing a polygenic trait with redundancy among loci that allows for a full range of adaptive architectures: from sweeps to subtle frequency shifts.

### 6.1 Polygenic architectures of adaptation

For any polygenic trait, the multitude of possible adaptive architectures is fully captured by the joint distribution of mutant alleles across the loci in its basis. Different adaptive scenarios (such as sweeps or shifts) correspond to characteristic differences in the shape of this distribution, at the end of the adaptive phase. For a single locus, the stationary distribution under mutation, selection, and drift can be derived from diffusion theory and has been known since the early days of population genetics (S. Wright (1931), [32]). For multiple interacting loci, however, this is usually not possible. To address this problem for our model, we dissect the adaptive process into two phases. The early stochastic phase describes the establishment of all mutants that contribute to the adaptive response under the influence of mutation and drift. We use that loci can be treated as independent during this phase to derive a joint distribution for ratios of allele frequencies at different loci, Eq (5). During the second, deterministic phase, epistasis and linkage become noticeable, but mutation and drift can be ignored. Allele frequency changes during this phase can be described as a density transformation of the joint distribution. For the simple model with fully redundant loci, and assuming either LE or complete linkage, this transformation can be worked out explicitly. Our main result Eq (8) can be understood as a multi-locus extension of Wright’s formula. For a neutral locus with multiple alleles, Wright’s distribution is a Dirichlet distribution, which is reproduced in our model for the case of complete linkage, see Appendix A.1. For the opposite case of linkage equilibrium, we obtain a family of inverted Dirichlet distributions, depending on the stopping condition – our time of observation.

Note that (unlike Wright’s distribution) the distribution of adaptive architectures is *not* a stationary distribution, but necessarily transient. It describes the pattern of mutant alleles at the end of the “rapid adaptive phase” [30, 31], because this is the time scale that the opposite narratives of population genetics and quantitative genetics refer to. In particular, the quantitative genetic “small shifts” view of adaptation does not talk about a stationary distribution: it does not imply that alleles will never fix over much longer time scales, due to drift and weak selection. On a technical level, the transient nature of our result means that it reflects the effects of genetic drift only during the early phase of adaptation. These early effects are crucial because they are magnified by the action of positive selection. In contrast, our result ignores drift after phenotypic adaptation has been accomplished – which is also a reason why it can be derived at all.

To capture the key characteristics of the adaptive architecture, we dissect the joint distribution in Eq (8) into marginal distributions of single loci. As explained at the start of the results section, these loci do not refer to a fixed genome position, but are defined *a posteriori* via their role in the adaptive process. For example, the *major locus* is defined as the locus with the highest mutant allele frequency at the end of the adaptive phase. (Since all loci have equal effects in our model, this is also the locus with the largest contribution to the adaptive response, but see Appendix A.4.) This is a different way to summarize the joint distribution than used in some of the previous literature [26, 28, 29], which rely on a gene-centered view to study the pattern at a focal locus, irrespective of its role in trait adaptation. In contrast, we use a trait-centered view, which is better suited to describe and distinguish adaptive scenarios. For example, “adaptation by sweeps” refers to a scenario where sweeps happen at some loci, rather than at a specific locus. This point is further discussed in Appendix A.6, where we also display marginal distributions of Eq (8) for fixed loci.

#### The role of the background mutation rate

Our results show that the qualitative pattern of polygenic adaptation is predicted by a single compound parameter: the background mutation rate Θ_*bg*_ (see Eqs (10),(13),(15)), i.e., the population mutation rate for the background of a focal locus within the trait basis. For a large basis, Θ_*bg*_ is closely related to the trait mutation rate. We can understand the key role of this parameter as follows. As detailed in the Section 4.4, the early stochastic phase of adaptation is governed by two processes: New successful mutations (destined for establishment) enter the population at rate Θ*_l_s_b_* per locus (where Θ_*l*_ is the locus mutation rate and *s_b_* the selection coefficient), while existing mutants spread with an exponential rate *s_b_*. Consider the locus that carries the first successful mutant. For Θ_*bg*_ < 1, the expected spread from this first mutant exceeds the creation of new mutant lineages at all other loci. Therefore, the locus will likely maintain its lead, with an exponentially growing gap to the second largest locus. Vice versa, for Θ_*bg*_ > 1, most likely one of the competing loci will catch up. We can thus think of Θ_*bg*_ as a measure of competition experienced by the major locus due to adaptation at redundant loci in its genetic background. The argument does not depend on the strength of selection, which affects both rates in the same way. The same can be shown for adaptation from standing genetic variation at mutation-selection-drift balance, see Mathematical Appendix (M.1). As a consequence of low mutant frequencies during the stochastic phase, the result is also independent of interaction effects due to epistasis or linkage.

Since the order of loci is not affected by the deterministic phase of the adaptive process, Θ_*bg*_ maintains its key role for the adaptive architecture. In the joint frequency distribution, Eq (5) and Eq (8), it governs the singular behavior of the marginal distribution at the major locus. For Θ_*bg*_ < 1, this distribution has a singularity at the maximum of its range. Adaptation is therefore dominated by the major locus, leading to heterogeneous architectures. For Θ_*bg*_ ≲ 0.1, adaptation occurs almost always due to a completed sweep at this locus. For Θ_*bg*_ > 1, in contrast, no single dominating locus exists: adaptation is collective and supported by multiple loci. For a polygenic trait with Θ_*bg*_ ≳ 100, we obtain homogeneous small shifts at many loci, as predicted by quantitative genetics.

The result also shows that the adaptive scenario does not depend directly on the number of loci in the genetic basis of the trait, but rather on their combined mutation rate (the mutational target size, *sensu* [11]). For redundant loci and fixed Θ_*bg*_, the predicted architecture at the loci with the largest contribution to the adaptive response is almost independent of the number of loci, see Fig 4. Qualitatively, the same still holds true when the assumption of complete redundancy is dropped (Fig 5). In this case, only loci in the genetic background that are not required to reach the new trait optimum, but offer redundant routes for adaptation, are included in Θ_*bg*_. Note that the same reasoning holds for a quantitative trait that is composed of several modules of mutually redundant genes, but where interactions among genes in different modules only affect a focal module as a unit. I.e., due to changes in the genetic background, all loci in this module experience a uniform change in the selection coefficient *s_b_* = *s_b_*(*t*) > 0. In this case, assuming LE, our model still applies (cf. the Mathematical Appendix). The adaptive architecture for each module depends only on the module-specific Θ_*bg*_, but not on the mutation rates at genes in the basis of the trait outside of the module.

#### Polygenic adaptation and soft sweeps

In our analysis of polygenic adaptation, we have not studied the probability that adaptation at single loci could involve more than a single mutational origin and thus produces a so-called *soft selective sweep from recurrent mutation*. As explained in [6, 40], however, the answer is simple and only depends on the locus mutation rate – independently of adaptation at other loci. Soft sweeps become relevant for Θ_*l*_ ≳ 0.1. For much larger values Θ_*l*_ ≫ 1, they become “super-soft” in the sense that single sweep haplotypes do not reach high frequencies because there are so many independent origins of the mutant allele. The role of Θ_*bg*_ for polygenic adaptation is essentially parallel to the one of Θ_*l*_ for soft sweeps. In both cases, the population mutation rate is the only relevant parameter, with a lower threshold of Θ ~ 0.1 for a signal involving multiple alleles and much higher values for a “super-soft” scenario with only subtle frequency shifts. Nevertheless, the mathematical methods to analyze both cases are different, essentially because the polygenic scenario does not lend itself to a coalescent approach.

### 6.2 Alternative approaches to polygenic adaptation

The theme of “competition of a single locus with its background” relates to previous findings by Chevin and Hospital (2008) [26] in one of the first studies to address polygenic footprints. These authors rely on a deterministic model of an additive quantitative trait to describe the adaptive trajectory at a single target QTL in the presence of background variation. The background is modeled as a normal distribution with a mean that can respond to selection, but with constant variance. Obviously, a drift-related parameter, such as Θ_*bg*_, has no place in such a framework. Still, there are several correspondences to our result on a qualitative level. Specifically, a sweep at the focal locus is prohibited under two conditions. First, the background variation (generated by recurrent mutation in our model, constant in [26]) must be large. Second, the fitness function must exhibit strong negative epistasis that allows for alternative ways to reach the trait optimum – and thus produces redundancy (due to Gaussian stabilizing selection in [26]). Finally, while the adaptive trajectory depends on the *shape* of the fitness function, Chevin and Hospital note that it does not depend on the *strength* of selection on the trait, as also found for our model.

A major difference of the approach used in [26] is the gene-centered view that is applied there. Consider a scenario where the genetic background “wins” against the focal QTL and precludes it from sweeping. For a generic polygenic trait (and for our model) this still leaves the possibility of a sweep at one of the background loci. However, this is not possible in [26], where all background loci are summarized as a sea of small-effect loci with constant genetic variance.

This constraint is avoided in the approach by deVladar and Barton [41] and Jain and Stephan [31], who study an additive quantitative trait under stabilizing selection with binary loci (see also [42] for an extension to adaptation to a moving optimum). These models allow for different locus effects, but ignore genetic drift. Before the environmental change, all allele frequencies are assumed to be in mutation-selection balance, with equilibrium values derived in [41]. At the environmental change, the trait optimum jumps to a new value and alleles at all loci respond by large or small changes in the allele frequencies. Overall, [41] and [31] predict adaptation by small frequency shifts in larger parts of the biological parameter space relative to our model. In particular, sweeps are prevented in [31] if most loci have a small effect and are therefore under weak selection prior to the environmental change. This contrasts to our model, where the predicted architecture of adaptation is independent of the selection strength. Thus, in our model, weak selection does not imply *shifts*. This difference can at least partially be explained by the neglect of drift effects on the starting allele frequencies in the deterministic models. In the absence of drift, loci under weak selection start out from frequency *x*_0_ = 0.5 [41]. In finite populations, however, almost all of these alleles start from very low (or very high) frequencies – unless the population mutation parameter is large (many alleles at intermediate frequencies at competing background loci are expected only if Θ_*bg*_ ≫ 1, in accordance with our criterion for *shifts*). To test this further, we have analyzed our model for the case of starting allele frequencies set to the deterministic values of mutation-selection balance, *μ*/*s_d_*. Indeed, we observe adaptation due to small frequency shifts in a much larger parameter range (Appendix A.2).

Generally, adaptation by sweeps in a polygenic model requires a mechanism to create heterogeneity among loci. This mechanism is entirely different in both modeling frameworks. While heterogeneity is (only) produced by unequal locus effects for the deterministic quantitative trait, it is (solely) due to genetic drift for the redundant trait model. Since both approaches ignore one of these factors, both results should rather underestimate the prevalence of sweeps. Indeed, heterogeneity increases for our model with unequal locus effects (see Appendix A.4).

Both drift and unequal locus effects are included in the simulation studies by Pavlidis et al (2012) [28] and Wollstein and Stephan (2014) [29]. These authors assess patterns of adaptation for a quantitative trait under stabilizing selection with up to eight diploid loci. However, due to differences in concepts and definitions there are few comparable results. In contrast to [31] and to our approach, they study long-term adaptation (they simulate *N_e_* generations). In [28, 29], *sweeps* are defined as fixation of the mutant allele at a focal locus, whereas *frequency shifts* correspond to long-term stable polymorphic equilibria [29]. With this definition, a *shift* scenario is no longer a transient pattern, but depends entirely on the existence (and range of attraction) of polymorphic equilibria. A polymorphic outcome is likely for a two-locus model with full symmetry, where the double heterozygote has the highest fitness. For more than two loci, the probability of shifts *decreases* (because polymorphic equilibria become less likely, see [43]). However, also the probability of a sweep decreases. This is largely due to the gene-centered view in [28], where potential sweeps at background loci are not recorded (see also Appendix A.6).

### 6.3 Scope of the model and the analytical approach

We have described scenarios of adaptation for a simple model of a polygenic trait. This model allows for an arbitrary number of loci with variable mutation rates, haploids and diploids, linkage, time-dependent selection, new mutations and standing genetic variation, and alternative starting conditions for the mutant alleles. Its genetic architecture, however, is strongly restricted by our assumption of (full or relaxed) redundancy among loci. In the haploid, fully redundant version, the phenotype is binary and only allows for two states, *ancestral wildtype* and *mutant*. Biologically, this may be thought of as a simple model for traits like pathogen or antibiotic resistance, body color, or the ability to use a certain substrate [44, 45].

Our main motivation, however, has been to construct a minimal model with a polygenic architecture that allows for both sweep and shifts scenarios – and for comprehensive analytical treatment. One may wonder how our methods and results generalize if we move beyond our model assumptions.

Key to our analytical method is the dissection of the adaptive process into a stochastic phase that explains the origin and establishment of beneficial variants and a deterministic phase that describes the allele frequency changes of the established mutant copies. This framework can be applied to a much broader class of models. Indeed, in many cases, the fate of beneficial alleles, establishment or loss, is decided while these alleles are rare. Excluding complex scenarios such as passage through a fitness valley, the initial stochastic phase is relatively insensitive to interactions via epistasis or linkage. We can therefore describe the dynamics of traits with a different architecture (e.g. an additive quantitative trait with equal-effect loci under stabilizing selection) within the same framework by coupling the same stochastic dynamics to a different set of differential equations describing the dynamics during the deterministic phase.

This is important because, as described above, the key *qualitative* results to distinguish broad categories of adaptive scenarios are due to the initial stochastic phase. This holds true, in particular, for the role of the background mutation rate Θ_*bg*_. We therefore expect that these results generalize beyond our basic model. Indeed, we have already seen this for our model extensions to include diploids, linkage, and relaxed redundancy. Vice-versa, we have seen that other factors, such as alternative starting conditions for the mutant alleles, directly affect the early stochastic phase and lead to larger changes in the results. As shown in Appendix A.2, however, they can be captured by an appropriate extension of the stochastic Yule process framework.

Several factors of biological importance are not covered by our current approach. Most importantly, this includes loci with different effect sizes and spatial population structure. Both require a further extension of our framework for the early stochastic phase of adaptation. Unequal locus effects (both directly on the trait or on fitness due to pleiotropy) are expected to enhance the heterogeneity in the adaptive response among loci, as confirmed by simulations of a 2-locus model in Appendix A.4. The opposite is true for spatial structure, as further discussed below.

### 6.4 When to expect sweeps or shifts

Although our assumptions on the genetic architecture of the trait (complete redundancy and equal loci) are favorable for a collective, shift-type adaptation scenario, we observe large changes in mutant allele frequencies (completed or partial sweeps) for major parts of the parameter range. A homogeneous pattern of *subtle frequency shifts* at many loci is only observed for high mutation rates. This contrasts with experience gained from breeding and modern findings from genome-wide association studies, which are strongly suggestive of an important role for small shifts with contributions from very many loci (reviewed in [1, 15, 46–48], see [12, 49, 50] for recent empirical examples). For traits such as human height, there has even been a case made for *omnigenic* adaptation [8], setting up a “mechanistic narrative” for Fisher’s (conceptual) infinitesimal model. Clearly, body height may be an extreme case and the adaptive scenario will strongly depend on the type of trait under consideration. Still, the question arises whether and how wide-spread shift-type adaptation can be reconciled with our predictions. We will first discuss this question within the scope of our model and then turn to factors beyond our model assumptions.

#### The size of the background mutation rate

The decisive parameter to predict the adaptive scenario in our model, the background mutation rate, is not easily amenable to measurement. Θ_bg_ = (*L* − 1)Θ_*l*_ compounds two factors, the locus mutation parameter Θ_*l*_ and the number of loci *L*, which are both complex themselves and require interpretation. To assess the plausibility of values of the order of Θ_*bg*_ ≳ 100, required for homogeneous polygenic shifts in our model, we consider both factors separately.

Large locus mutation rates Θ_*l*_ = 4*N_e_μ* (for diploids, 2*N_e_μ* for haploids) are possible if either the allelic mutation rate *μ* or the effective population size *N_e_* is large. Both cases are discussed in detail (for the case of soft sweeps) in [6]. Basically, *μ* can be large if the mutational target *at the locus* is large. Examples are loss-of-function mutations or cis-regulatory mutations. *N_e_* is the *short-term effective population size* [40, 51, 52] during the stochastic phase of adaptation. This *short-term* size is unaffected by demographic events, such as bottlenecks, prior to adaptation. It is therefore often larger than the *long-term* effective size that is estimated from nucleotide diversity. (Strong changes in population size *during* the adaptive period can have more subtle effects [53].) For recent adaptations due to gain-of-function mutations, plausible values are Θ_*l*_ ≲ 0.1 for *Drosophila* and Θ_*l*_ ≲ 0.01 for humans [6].

If 10 000 loci or more contribute to the basis of a polygenic trait [8], large values of Θ_*bg*_ could, in principle, easily be obtained. However, the parameter *L* in our model counts only loci that actually can respond to the selection pressure: mutant alleles must change the trait in the right direction and should not be constrained by pleiotropic effects. Omnigenic genetics, in particular, also implies ubiquitous pleiotropy and so the size of the basis *that is potentially available for adaptation* is probably strongly restricted. For a given trait, the number of available loci *L* may well differ, depending on the selection pressure and pleiotropic constraints. Furthermore, our results for the model with relaxed redundancy show that Θ_*bg*_ only accounts for loci that are truly redundant and offer alternative routes to the optimal phenotype. With this in mind, values of *L* in the hundreds or thousands (required for Θ_*bg*_ ≥ 100) seem to be quite large. While some highly polygenic traits such as body size could still fulfill this condition, this appears questionable for the generic case.

#### Balancing selection and spatial structure

In our model, characteristic patterns in the adaptive architecture result from heterogeneities among loci that are created by mutation and drift during the initial stochastic phase of adaptation. As initial condition, we have mostly assumed that mutant alleles segregate in the population in the balance of mutation, purifying selection and genetic drift. Since this typically results in a broad allele frequency distribution (unless mutation is very strong), it favors heterogeneity among loci and thus adaptation by (partial) sweeps. However, even after decades of research, the mechanisms to maintain genetic variation in natural populations remain elusive [1]. As discussed in Appendix A.2, more homogeneous starting conditions for the mutant alleles can be strongly favorable of a shift scenario. Such conditions can be created either by balancing selection or by spatial population structure.

Balancing selection (due to overdominance or negative frequency dependence) typically maintains genetic variation at intermediate frequencies. If a major part of the genetic variance for the trait is due to balancing selection, adaptation could naturally occur by small shifts. However, the flexibility of alleles at single loci, and thus the potential for smaller or larger shifts, will depend on the strength of the fitness trade-off (e.g. due to pleiotropy) at each locus. If these trade-offs are heterogeneous, the adaptive architecture will reflect this. Also, adaptation against a trade-off necessarily involves a fitness cost. Therefore, if the trait can also adapt at loci that are free of a trade-off, these will be preferred, possibly leading to sweeps.

As discussed in a series of papers by Ralph and Coop [34, 35], spatial population structure is a potent force to increase the number of alternative alleles that contribute to the adaptive response. If adaptation proceeds independently, but in parallel, in spatially separated subpopulations, different alleles may be picked up in different regions. Depending on details of the migration pattern [36], we then expect architectures that are globally polygenic with small shifts, but locally still show sweeps or dominating variants.

Furthermore, population structure and gene flow *before* the start of the selective phase can have a strong effect on the starting frequencies. In particular, if the base population is admixed, mutant alleles could often start from intermediate frequencies and naturally produce small shifts. This applies, in particular, to adaptation in modern human populations, which have experienced major admixture events in their history [54, 55] and only show few clear signals of selective sweeps [11].

Finally, gene flow and drift will continue to change the architecture of adaptation after the rapid adaptive phase that has been our focus here. This can work in both directions. On the one hand, subsequent gene flow can erase any *local* sweep signals by mixing variants that have been picked up in different regions [34, 35]. On the other hand, local adaptation, in particular, may favor adaptation by large-effect alleles at few loci, favoring sweeps over longer time-scales. Indeed, as argued by Yeaman [56], initial rapid adaptation due to small shifts at many alleles of mostly small effect may be followed by a phase of allelic turnover, during which alleles with small effect are swamped and few large-effect alleles eventually take over. This type of allele sorting over longer time-scales is also observed in simulations studies for a quantitative trait under stabilizing selection that adapts to a new optimum after an environmental change [31, 57].

#### Between sweeps and shifts: adaptation by partial sweeps

Previous research has almost entirely focused on either of the two extreme scenarios for adaptation: sweeps in a single-locus setting or (infinitesimal) shifts in the tradition of Fisher’s infinitesimal model. This leaves considerable room for intermediate patterns. Our results for the redundant trait model show that such transitional patterns should be expected in a large and biologically relevant parameter range (values of Θ_*bg*_ between 0.1 and 100). Patterns between sweeps and shifts are *polygenic* in the sense that they result from the *concerted* change in the allele frequency at multiple loci. They can only be understood in the context of interactions among these loci. However, they usually do not show subtle shifts, but much larger changes (partial sweeps) at several loci. If adaptation occurs from mutation-selection-drift balance, the polygenic patterns are typically strongly heterogeneous, even across loci with identical effects on the trait. Such patterns may be difficult to detect with classical sweep scans, in particular if partial sweeps are “soft” because they originate from standing genetic variation or involve multiple mutational origins. However, they should be visible in time-series data and may also leave detectable signals in local haplotype blocks.

Indeed there is empirical evidence for partial sweeps from time series data in experimental *evolve and resequence* experiments on recombining species such as fruit flies. For example, Burke *et al*. [58] observe predominantly partial sweeps (from SGV) in their long-term selection experiments with *Drosophila melanogaster* for accelerated development – a rather unspecific trait with a presumably large genomic basis. A similar pattern of “plateauing”, where allele frequencies at several loci increase quickly over several generations, but then stop at intermediate levels, was recently observed by Barghi and collaborators [59] for adaptation of 10 *Drosophila simulans* replicates to a hot temperature environment. Complementing the genotypic time-series data with measurements of several phenotypes, these authors found convergent evolution for several high-level traits (such as fecundity and metabolic rate), indicating that rapid phenotypic adaptation had reached a new optimum. This high-level convergence contrasts a strong heterogeneity in the adaptation response among loci and also between replicates [59]. Based on their data, the authors reject both a selective sweep model and adaptation by subtle shifts. Instead, the observed patterns are most consistent with the intermediate adaptive scenario in our framework, featuring heterogeneous partial sweeps at interacting loci with a high level of genetic redundancy.

## A Supporting information

### A.1 Linked loci

Negative epistasis for fitness causes negative linkage disequilibrium (LD) among the selected loci. While LD can typically be ignored as long as loci are only loosely linked, this changes once recombination rates drop below a threshold (e.g. [22], p. 277). For tight linkage *r* → 0, in particular, individuals carrying multiple mutations can no longer be formed by recombination, but require multiple mutational hits on the same haplotype. This is unlikely while mutant allele frequencies are low, which is when the relevant mutations of the adaptive process arise. By the end of the adaptive phase, the excess of single-mutant haplotypes produces strong negative LD. Nevertheless, our theory predicts that the distribution of allele frequency ratios that emerges from the early stochastic phase of the adaptive process is unaffected Eq.(9). This prediction is confirmed by simulations, see Fig S.1.

Fig S.2 shows the joint distribution of the major and the minor locus of a trait with *L* = 2 loci for different degrees of linkage. In all cases, the process is stopped when the proportion of remaining non-mutant individuals drops below *f_w_* = 0.05. The results show that the linkage equilibrium assumption (red and blue lines) provides a good approximation as long as *r* ≥ *s_b_*. For *r* < *s_b_*, the distributions are shifted to lower values and clear deviations become visible. The constraint on the allele frequencies at the stopping condition changes from (1 − *p*_1_)(1 − *p*_2_) = *f_w_* for linkage equilibrium to *p*_1_ + *p*_2_ = 1 − *f_w_* for complete linkage. As a consequence, the boundary between the major and minor locus distributions (red and blue) drops from 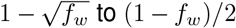. As shown in the Mathematical Appendix, Eq (M.29), we can derive an analytical approximation for the distributions for complete linkage *r* = 0. For *L* = 2, we obtain a modified Beta-distribution (black lines in the Figure)

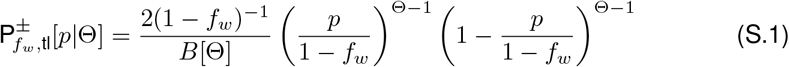

with *p* ≥ (1 − *f_w_*)/2 (resp. *p* ≤ (1 − *f_w_*)/2) for the major (minor) locus. The simulation 1 results show that this prediction is accurate for *r* ≪ *s_b_* (deviations for Θ_*bg*_ = 100 are due 1 to overshooting of the stopping condition in the last generation of our Wright-Fisher simulations).

While linkage affects the shape of the joint distribution, it does not alter its key qualitative characteristics that distinguish adaptive scenarios. In particular, the same conditions on Θ_*bg*_ and Θ_*l*_ apply for singularities at the boundaries of the marginal distributions. We still observe sweep-like adaptation for Θ_*bg*_ ≪ 1, adaptation by small shifts for Θ_*bg*_ ≫ 1, and a heterogeneous pattern of partial sweeps in a transition range of Θ_*bg*_ around 1.

**Fig S.1.**
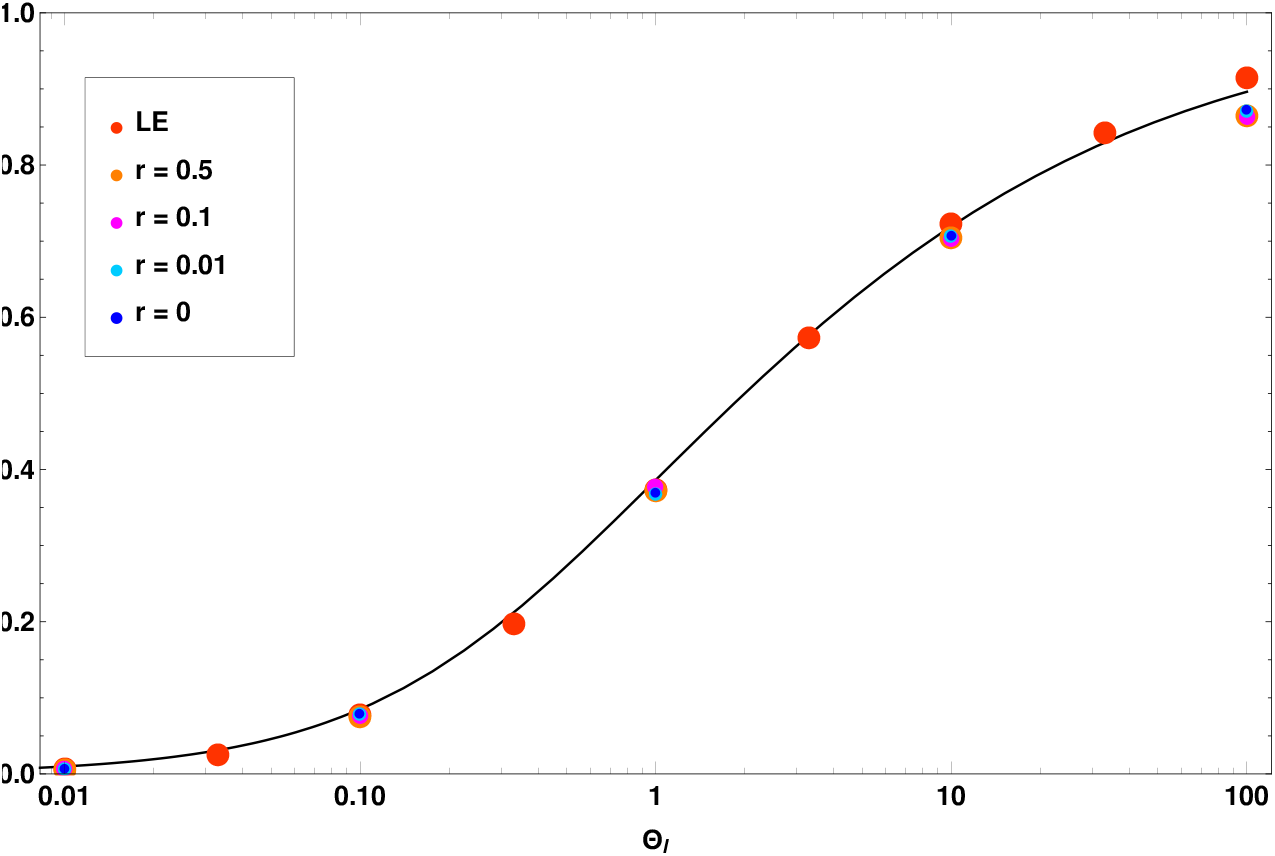
E[*x*] for redundant fitness effects with two linked loci. Simulation results (colored dots) for the mean allele frequency ratio are plotted in dependence of the locus population mutation rate Θ_*l*_ and compared with the analytical prediction (black line). Simulations are stopped when fitness has reached 95% of its maximum. Linkage does not change the results for the ratio of allele frequencies, despite significant buildup of linkage disequilibrium with low recombination rates. Results for 10 000 replicates, standard errors < 0.005 (smaller than symbols).

### A.2 Alternative starting allele frequencies

So far, we have assumed that adaptation starts from mutation-selection-drift balance. This includes variable amounts of standing genetic variation (weak or strong *s_d_*) and even cases where this balance is not represented by a stable equilibrium distribution (time-dependent selection, see the Mathematical Appendix). There are, however, other scenarios of biological relevance. Given the right (possibly complex) selection scheme, balancing selection can maintain mutant alleles, prior to the environmental change, at arbitrary frequencies. The same holds true if the base population is admixed, either due to natural processes or due to human activity (e.g. breeding from hybrids). For these scenarios, our theoretical formalism to describe the establishment of mutants during the stochastic phase (Fig 2) does not apply. In this section, we describe how the formalism can be extended to cover arbitrary starting frequencies of mutants at the onset of positive selection at time *t* = 0.

**Fig S.2.**
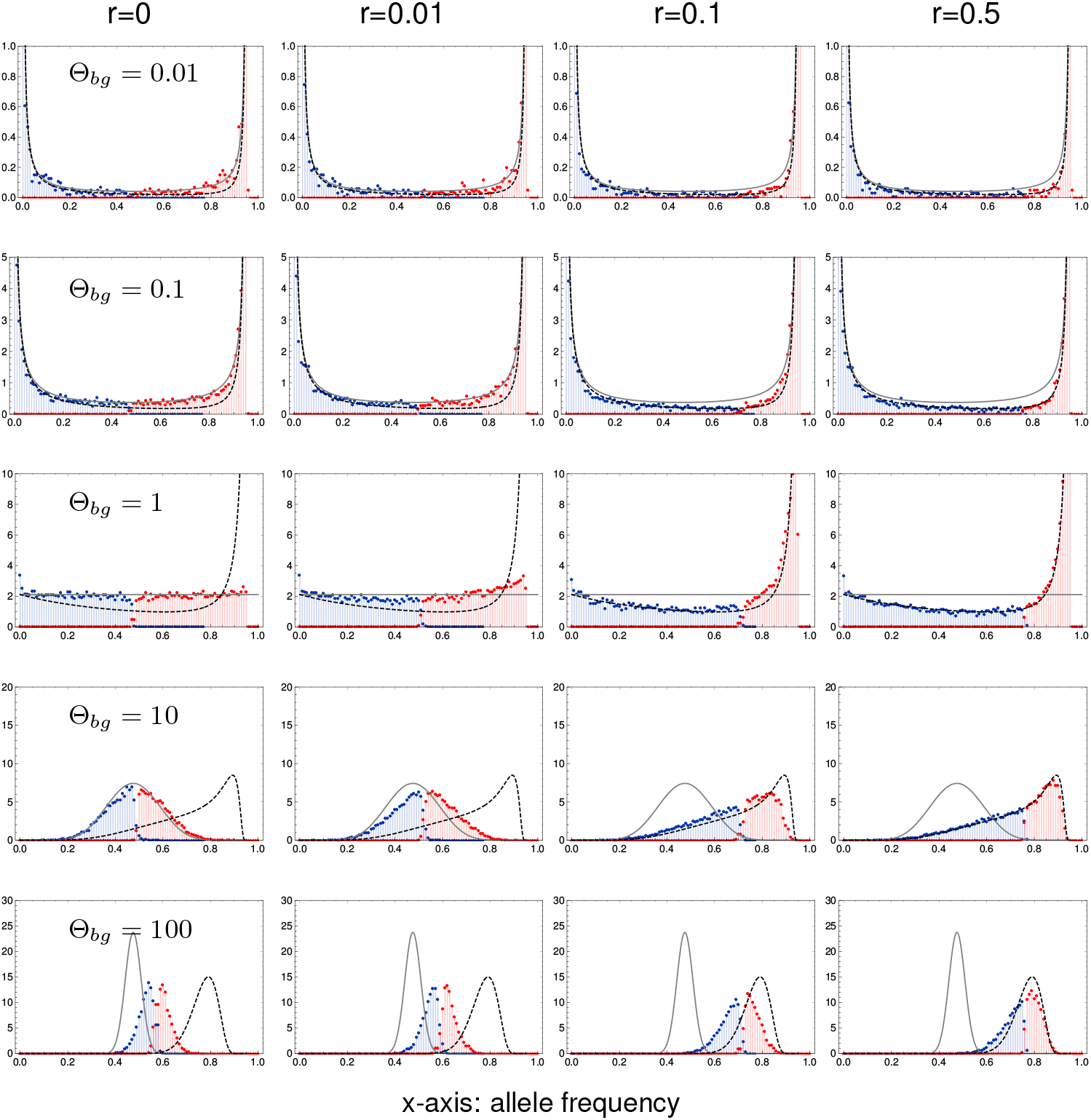
Genetic architecture of adaptation with linkage. Marginal distributions for the major locus (red) and the minor locus (blue) of a model with *L* = 2 loci depending on Θ_*bg*_ (rows) and linkage among the loci(columns). Black lines show the analytical approximations for LE (dashed) and complete linkage (solid). For strong recombination *r* ≥ *s_b_* = 0.1, the deviations from the LE approximation are small. For *r* ≪ *s_b_* = 0.1, the approximation for complete linkage works well. Further parameters: −*s_d_* = *s_b_* = 0.1, *N_e_* = 10 000,10 000 replicates.

#### Extended Yule framework

The Yule process that describes the stochastic phase of the adaptive process accounts for the mutant copies at all loci that are destined for establishment. In our framework so far (see the Mathematical Appendix M.2), we have started this process with zero copies. SGV due to mutation-selection-drift balance can still be produced by such a process if it is started at some time in the past (*t* < 0). For general starting frequencies, we can alternatively start this process at time *t* = 0, but with mutant copies (immortal lineages) already present. Suppose that the mutant frequency at locus *i* at time *t* = 0 is *p_i_*, corresponding to *N_e_p_i_* mutant copies. Of these, only the *n_i_* < *N_e_p_i_* “immortal” mutants (destined for establishment) are included in the Yule process. Assuming an independent establishment probability *p*_est_ per copy, *n_i_* is binomially distributed with parameters *N_e_p_i_* and *p*_est_. For the limit distribution of a multi-type Yule process that is started with a non-zero number of lines, consider that each of these initial lines can be understood as an extra source of new immortal lines (due to birth) that is entirely equivalent to the generation of new lineages by mutation. It is therefore appropriate to include these lines as *extra locus mutation rate*

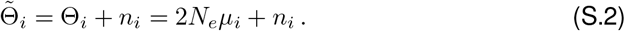

In the absence of recurrent mutation, Θ_*i*_ = 0, this procedure reproduces the well-known Polya urn scheme (e.g. [60, 61]). Replacing Θ_*i*_ by 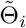 within our original Yule process formalism, and averaging over the binomial distribution, leads to the desired extension to arbitrary starting frequencies.

#### Application

Theory papers (e.g. [30, 31, 41, 62]) often use a deterministic framework to describe the frequency of alleles that segregate in a population in mutation-selection balance. To simplify the analysis, they do not model SGV as a distribution (due to mutation, selection, and drift), but replace this distribution by its expected value (ignoring drift). We can apply our scheme with fixed starting frequencies to this case and thus assess the effect of genetic drift in the starting allele frequency distribution. We assume equal loci and a starting frequency |*μ_l_*/*s_d_*| for an (initially deleterious) mutant allele with selection coefficient *s_d_* in mutation-selection balance. Fig S.3 shows the simulated marginal distributions of the loci with the largest contribution to the adaptive response (compare Fig 4). We see that the type of the adaptive architecture is again constant across rows with equal background mutation rate. However, due to the more homogeneous starting conditions, adaptation involves more loci and is much more shift-like. Analytical predictions following the above scheme are shown for *L* = 2 loci. With establishment probability *p*_est_ = 2*s_b_*, the counts *n*_1_ and *n*_2_ of ‘‘immortal” mutants at both loci are independent random draws from a Binomial distribution with parameters *N_e_*|*μ_l_*/*s_d_*| = |Θ_*l*_/2*s_d_*| and 2*s_b_*. For Θ_*bg*_ ≥ 0.1, we find (heuristically) that the marginal distribution for alleles starting from mutation-selection balance closely matches the one of the fully stochastic model with effective 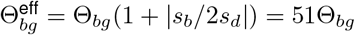 for the parameters in the figure (lines added in green). (Note that, from the average number of established lines, one would assume 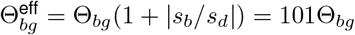. However, this does not account for the variance in the number of immortal lines among the two loci.)

### A.3 Diploids

To extend our model to diploids, we assume that a single locus that is *homozygous* for the mutant allele is sufficient to produce the fully functional mutant phenotype, while a *heterozygous* locus produces a mutant that is functional with probability 1 − *h*. We assume that mutants contribute independently. Thus, if *k* heterozygous loci exist, but no homozygous mutant locus, the resulting mutant phenotype will be functional with probability 1 − (1 − (1 − *h*))^*k*^ = 1 − *h^k^*. For *L* = 2 loci, in particular, the (logarithmic) fitness of genotype *G* becomes

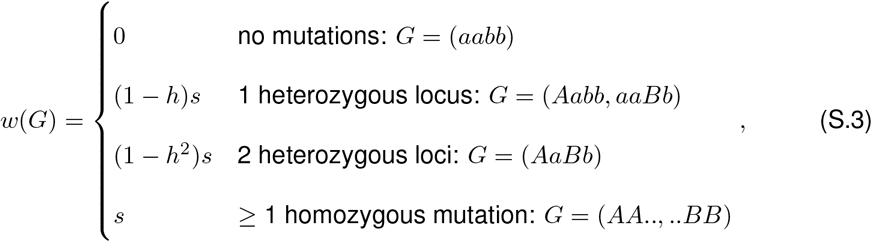

where *s* = *s_b_* > 0 for *t* ≥ 0 and *s* = *s_d_* < 0 for *t* < 0. Note that *h* ∈ [0, 1] measures the dominance of the *ancestral* allele. We assume Hardy-Weinberg-linkage-equilibrium (HWLE). In this case, the marginal fitnesses of the mutant alleles are (for 2 loci),

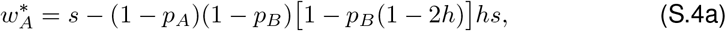

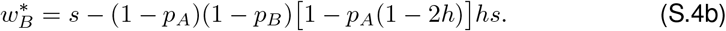

**Fig S.3.**
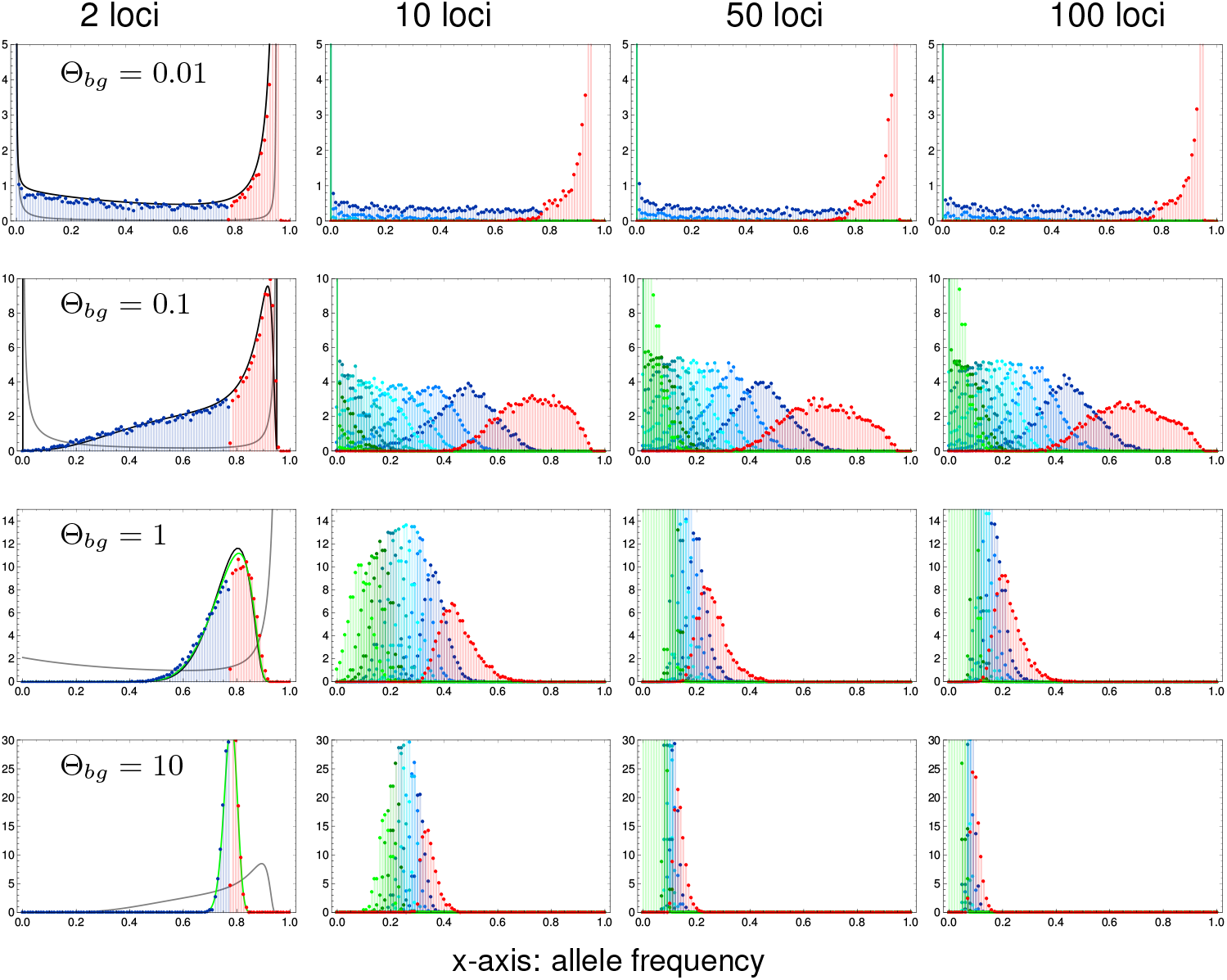
Polygenic adaptation from alternative allele starting frequencies. The panels show the adaptive architecture when mutant alleles start from their expected value in mutation-selection balance, without drift. We distribute *L* · |Θ_*l*_/2*s_d_*| mutant copies as evenly as possible across all loci. We set −*s_d_* = *s_b_*/100 = 0.001. Black lines for *L* = 2 loci show analytical predictions described in the main text (only computationally possible for Θ_*bg*_ ≤ 1), green lines for Θ_*bg*_ ≥ 1 show the heuristic prediction for 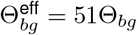. Finally, gray lines show the marginal distributions when adaptation occurs from mutation-selection-drift balance, compare Fig 4.

In contrast to the haploid case, the marginal fitnesses are in general *not* equal. There are, however, two important special cases, where our fitness scheme (with redundancy on the level of loci) implies equal marginal fitnesses (and thus redundancy on the level of alleles): either if the ancestral allele is fully recessive (*h* = 0) or if the alleles are co-dominant (*h* = 0.5). As shown in the Mathematical Appendix, this holds true more generally for an arbitrary number of loci.

#### Simulation results

We simulated a diploid model with two loci in HWLE according to the above scheme with three different levels of dominance of the ancestral allele, *h* = 0.1; 0.5; and 0.9. The diploid, effective population size is *N_e_*, corresponding to 2*N_e_* chromosomes. The mutation rate is *μ* at both loci and we define the population-scaled mutation rate for diploids as 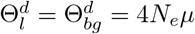. Simulations are stopped when the percentage of remaining ancestral *haplotypes* drops below *f_w_* = 0.05. (This condition directly corresponds to the stopping condition for haploids. Alternative stopping conditions, such as 95% increase in mean diploid fitness are also covered by our theoretical framework, but require a different transformation.)

The results are shown in Fig S.4. We see that the haploid results fully carry over to diploids for co-dominance (*h* = 0.5, middle column), where the diploid fitness scheme implies redundancy on the level of alleles. As explained above, the same holds true if the ancestral allele is fully recessive. Our simulations show that the haploid result is still a good approximation for *h* = 0.1 (left column). In contrast, much larger deviations are obtained for recessive mutants (dominant ancestral allele, *h* = 0.9, right column). In this case, the locus with the higher mutant frequency experiences stronger selection. For Θ_*l*_ ≥ 0.1, when polymorphism at both loci is likely, this favors the major locus relative to the minor locus, increasing the heterogeneity in the adaptive architecture.

### A.4 Asymmetric loci

For the Figures in the main text, we have assumed that all loci in the genetic basis of the trait are equivalent: they have equal mutation rates and effect sizes. This symmetric choice favors a collective *shift* scenario because no locus has a build-in advantage. In this Appendix, we study the consequences of asymmetries among loci.

#### Mutation rate asymmetry

Our analytical formalism allows for arbitrary asymmetries in the locus mutation rates. The prediction for the expected ratio of minor/major locus frequencies of a 2-locus model with unequal mutation rates Θ_1_ and Θ_2_ reads

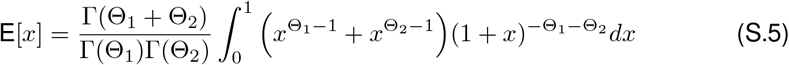

where the sum in the integral accounts for the possibility that either locus may end up as the “major locus” at the time of observation (compare Eq. M.27). Fig S.5 shows the prediction as a function of Θ_1_ and Θ_2_ = *d*Θ_1_ together with simulation results (analogous to Fig 3 in the main text). As expected, differences in the locus mutation rates lead to more heterogeneous “sweep-like” architectures with lower minor/major locus ratio. The Figure also confirms the independence of levels of standing genetic variation and the good overall fit of the analytical approximation.

**Fig S.4.**
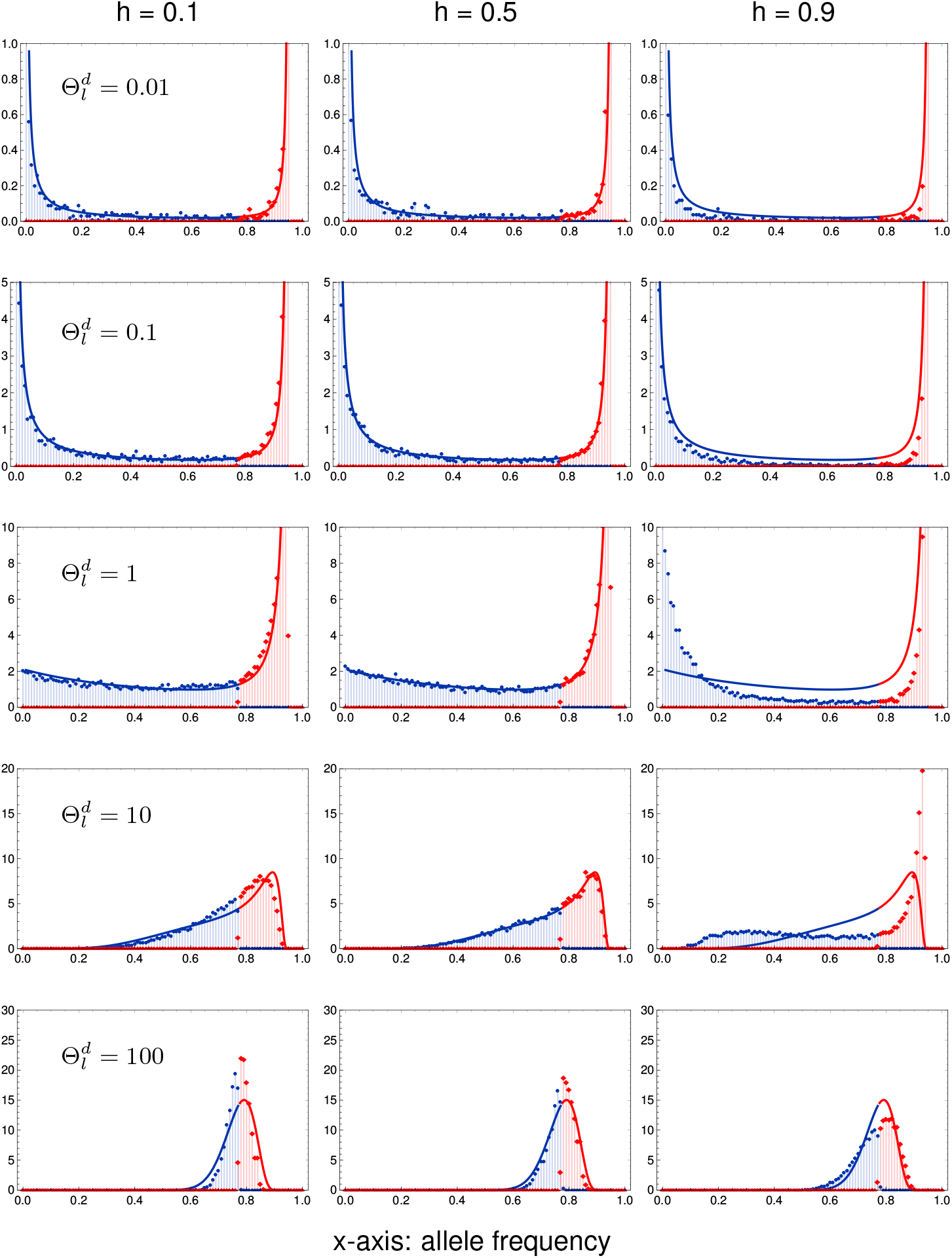
Adaptive architecture for diploids in linkage equilibrium. Adaptation in a 2-locus model according to scheme (S.3), with recessive (*h* = 0.1), codomiant (*h* = 0.5) or dominant (*h* = 0.9) ancestral alleles. We assume Hardy-Weinberg and linkage equilibrium. Simulations are stopped when frequency of wildtype haplotypes drops below 5%. Standing genetic variation builds up for 16*N_e_* generations before the change in the environment. Selection coefficients are set to *s_b_* = −*s_d_* = 0.1. Solid lines show analytical predictions using the framework developed for haploids.

**Fig S.5.**
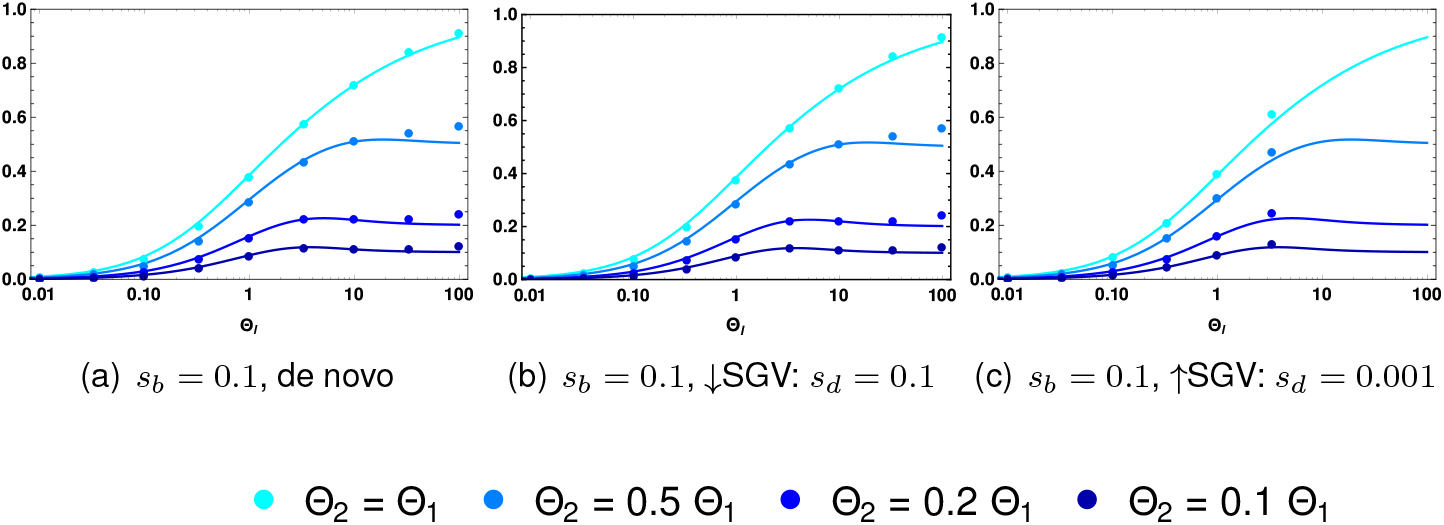
Different mutation rates. For *L* = 2 we plot E[*x*] without and with previous buildup of weak and strong SGV for different mutation rates at the two loci, such that Θ_2_ = *d* · Θ_1_, for *d* = 1, 0.5, 0.2, 0.1. Our analytical predictions for different mutation rates, Eq (S.5), yield an excellent fit. Simulations are obtained from 10 000 replicates per data point, assuming linkage equilibrium.

#### Locus effect asmmetry

Our analytical results are based on the assumption of strong redundancy between loci. In the main text, we have already discussed how these results extend for a scenario of relaxed redundancy, where two mutational steps are needed to reach the trait optimum. Similarly, intermediate phenotypes are also included in the diploid version of our model. However, both model extensions do not break the symmetry assumption concerning the effects of single-locus substitutions. Differences in the single-locus effects interfere with the assumptions of our Yule-process framework for the early adaptive phase. In contrast to unequal mutation rates, they cannot easily be included. Although polygenic models with equal locus effects have a long history in the biological literature, at least slight deviations from this assumption are unavoidable in nature. Indeed, deviations already arise due to non-neutral “hitchhiker” mutations on the selected haplotypes. With exponential growth during the selected phase, even small perturbations could, in principle, lead to significant changes in the resulting adaptive architecture. To test this, we use a haploid 2-locus model with (Malthusian) fitness 0 for the ancestral genotype *ab* and fitness *s*_*b*/*d*_ ≷ 0 for the single mutant *Ab* and the double mutant *AB*. The other single mutant, *aB* is set to *ϵs_b/d_*. Fig S.6 shows simulation results for the expected minor/major frequency ratio for cases where *aB* is less beneficial (*ϵ* = 100/101, 10/11, 2/3) as well as for cases where *aB* is optimal (*ϵ* = 101/100, 11/10, 3/2). Note that the latter case corresponds to “sign epistasis” for the *A* mutant. Simulations are stopped when the frequency of ancestral haplotypes, *ab*, drops below 5%.

As expected, the results show that unequal locus effects (like unequal mutation rates) lead to more heterogeneous adaptive architectures. However, as long as differences in the locus effects are moderate (below ~ 10%) the prediction from the fully redundant model still provides a good approximation. In contrast, differnces of 50% in the single-locus effects lead to sizable deviations. This relative robustness is reminiscent of the case of *soft selective sweeps*, where differences of ≲ 20% in the fitness of independent mutant copies only lead to small deviations from the predictions for the frequencies of sweep haplotypes (see Fig. 4 and S1 in [40]). Deviations from the fully redundant prediction are larger for the sign-epistasis case, where the aB mutant has the highest fitness. This is expected – indeed, the single mutant should eventually displace all other genotypes at later observation times. Fig S.6 also shows that deviations are partially compensated if adaptation occurs from standing genetic variation, in particular if levels of standing variation are high (panel c). This reflects our model assumption that the locus under stronger beneficial selection is also under stronger deleterious selection prior to the environmental shift.

**Fig S.6.**
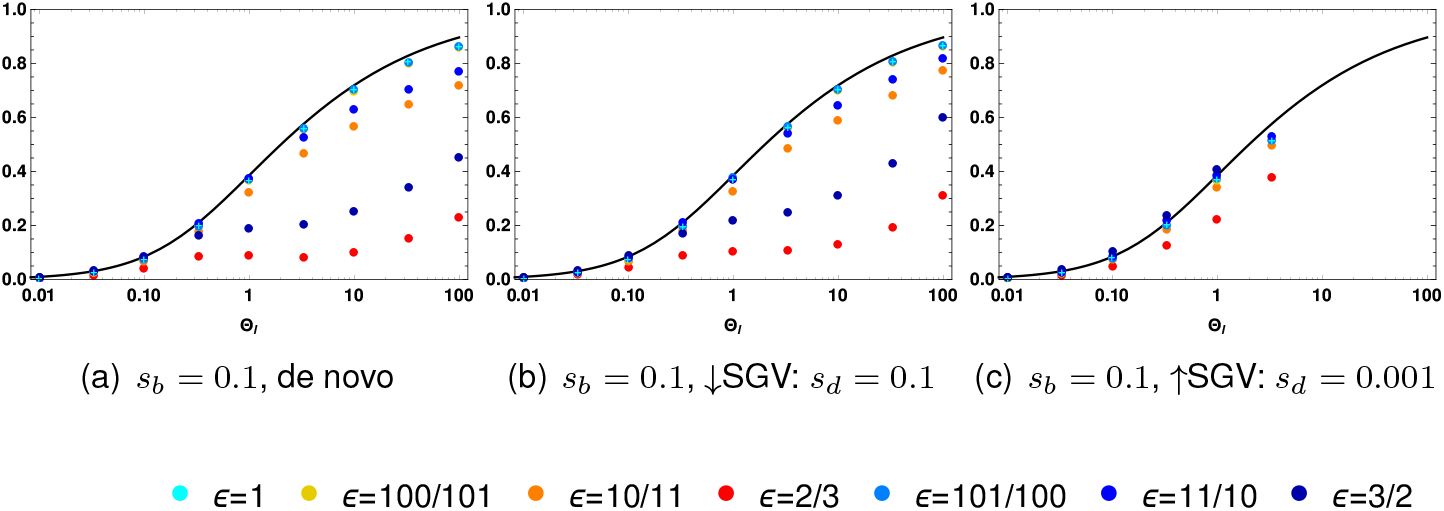
Different locus effects. For *L* = 2 we plot E[*x*] for without and with previous build-up of weak and strong SGV for various genotypic fitnesses of the aB-genotype *ϵs_b/d_*. Fitness of the Ab and AB genotype is always set to *s_b/d_*. Simulations are obtained from 10 000 replicates per mutation rate with recombination rate *r* = 0.5.

### A.5 Approximations for multi-locus architectures

For tight linkage, where the joint distribution of mutant alleles is given by a Dirichlet distribution, Mathematical Appendix Eq (M.29), lower dimensional marginal distributions for single locior groups of locican easily be derived. For linkage equilibrium, Mathematical Appendix Eq (M.20), however, the required integrals can only be solved numerically. For *L* loci, an (*L* – 2)-dim integral needs to be evaluated, which becomes computationally unfeasible (with programs packages like *Mathematica)* for *L* > 5. In many cases, we can nevertheless derive approximations. To do so, we make use of a key property of the adaptive architecture, seen in our results: The (joint) architecture of adaptation at loci with the largest contribution to the adaptive response is primarily a function of combined mutation rates at competing loci, such as the background mutation rate Θ_*bg*_. Given these values, it is largely independent of the number of loci in the genetic basis of the trait itself. We can therefore describe the adaptive architecture of a polygenic trait with L loci by a model with *k* < *L* loci *given that* the total adaptive response is well captured by the contribution of the top k loci. It turns out that this is typically the case for Θ_*bg*_ ≤ 1, when the contributions from different loci are very heterogeneous. In the following, we describe this procedure for an *L*-locus model with equal mutation rates Θ_*i*_ = Θ_*l*_, 1 ≤ *i* ≤ *L*.

#### Approximations using the 2-locus model

Several key properties of the *L*-locus architecture can already be described within the 2-locus framework. This includes the marginal distributions at the major locus and at the first minor locus. To this end, we set the mutation rate at the minor locus of the 2-locus model to the background mutation rate of the *L*-locus model. As described in the main text, this choice matches the time lag between the first origin of a mutation destined for establishment at a locus (usually the major locus) and at a second locus (usually the first minor locus). It also guarantees that the approximation captures the correct asymptotic shape of the major-locus distribution at *p* = 1 – *f_w_*, and of the first-minor-locus distribution at *p* = 0. The choice of the mutation rate at the major locus itself is less important. For the approximation of the major-locus distribution, we find that setting it to the locus-mutation rate yields the best fit. We thus use a 2-locus model with unequal mutation rates, 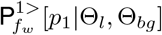, Eq (M.28a), in Fig 4. For the marginal distribution at the first minor locus, the approximation with equal mutation rates, 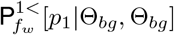, Eq (M.28b), works slightly better. Finally, we can also approximate the distribution at an *average* minor locus (rather than the first minor locus) by 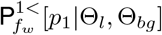.

#### Approximations using models with *k* ≥ 2 loci

The approximation of higher-order minor loci requires models with a sufficiently large genetic basis that such a locus exists at all. I.e., a *k*-locus model can approximate marginal distributions up to the (*k* – 1)st minor locus. Assume that we want to approximate the marginal distribution of the *j*th minor locus of an *L*-locus model using a *k*-locus model, *j* < *k* < *L*. As for the case *k* = 2 discussed above, the approximation requires that the expected lag time between the origin of a successful mutation at a first locus and the origin of a mutation at a *j*th locus be matched. For the *L*-locus model, this waiting time is

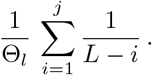

For a *k*-locus model with equal mutation rate 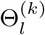 at all loci, we thus obtain the matching rule

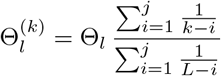

**Fig S.7.**
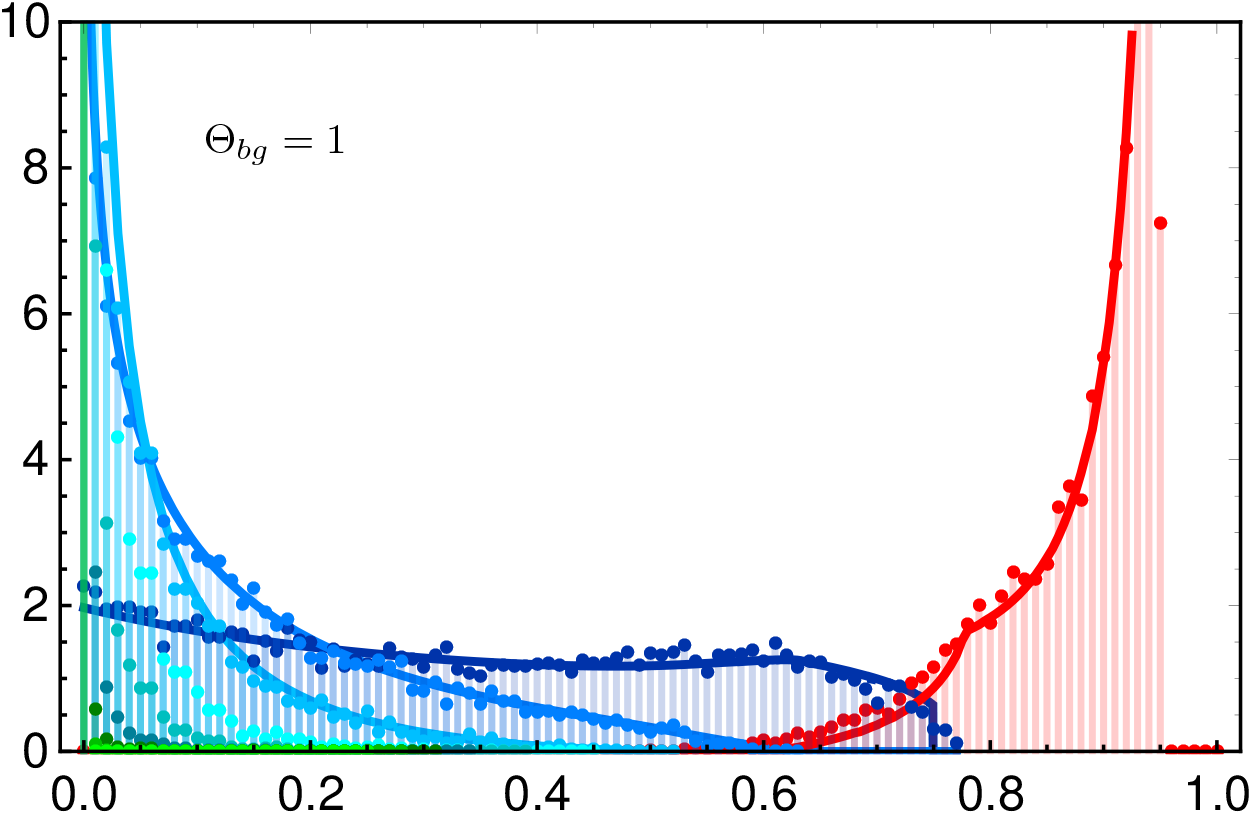
Approximating higher dimensional adaptive architectures. We approximate a 10 locus model (Θ_*bg*_ = 1) with the theoretical predictions based on the four-locus model for the major locus and the first, second, and third minor locus. Compare Fig 4, where we use approximations based on models with the minimal number of loci needed.

for the approximation of the jth minor locus. For *j* = 1, this reproduces the matching rule for the background mutation rate Θ_*bg*_. In general, the value for 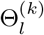 depends on *j*, but converges once *L, k* ≫ *j*. Approximations by models with unequal locus mutation rates are also possible, but usually do not lead to a relevant improvement. In Fig 4, we use formulas from 3- and 4-locus models to approximate the marginal distributions of the 2nd and 3rd minor locus, respectively. In general, the approximations for all loci can be improved by using approximation models with more loci than required, i.e. *k* > *j* + 1. In Fig S.7, we show this for approximations of the major locus and the first three minor loci, all derived from a 4-locus model.

### A.6 Marginal distribution of a single locus

Figure S.8 shows the marginal distribution at a single focal locus for a trait with *L* = 2 to *L* = 100 loci in its basis. Since all loci are equal, the probability that the focal locus ends up as the major locus is 1/*L*. The red dots in the figure indicate the part of the marginal distribution that corresponds to this case. With an increasing number of redundant loci, the probability for each single locus to play a major role in the adaptive process decreases. The marginal distribution of a fixed locus therefore changes strongly with an increasing number of loci *L*. For large *L*, in particular, it does not represents the key components of the adaptive architecture on the level of the trait any more. This is in contrast to Fig 4, where marginal distributions of the loci with the largest contributions to the adaptive response are shown. For 2 loci, Fig S.8 also shows the analytical approximation for the marginal distribution, Eq (11). As long as the adaptive architecture is dominated by only a few loci, the same 2-locus result can be used as an approximation for the marginal distribution in models with more than two loci. This is shown in the figure for Θ_*bg*_ ≤ 1. The figure also shows that the approximation fails for Θ_*bg*_ ≥ 10 when adaptation is truly collective.

**Fig S.8.**
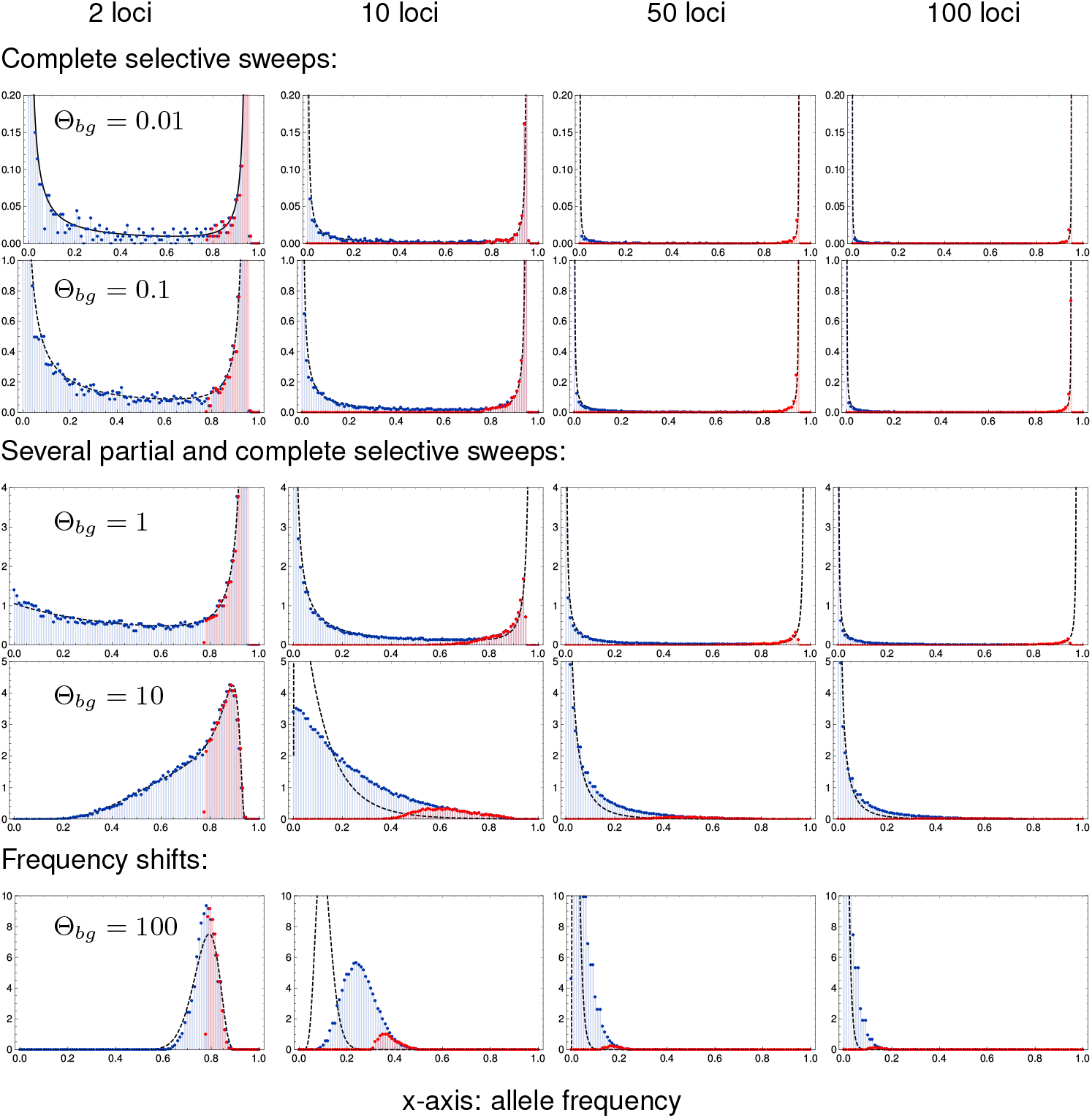
Marginal distribution at a single focal locus. Simulation results for the marginal distribution at a single locus at the end of the adaptive phase are shown in blue. Red dots show the contribution of the major locus to this distribution (all cases, where the focal locus ends up as the major locus). Dashed lines show the analytical prediction based on the 2-locus model, Eq (11). Parameters and further details as in Fig 4.

### A.7 Dynamics of adaptation

In contrast to previous work on the topic (e.g. [30,31]), our approach does not discuss adaptive architecture as a function of the time that has elapsed since the environmental change. Instead, we assess adaptation at the genotypic level as a function of the progress that has been made towards adaptation of the trait. In our main result on the joint distribution of mutant allele frequencies (Eq (8), this progress is measured by the stopping condition *f_w_*, which directly relates to the distance of the trait mean to the new optimum (see Eq (2); for the basic model of a fully redundant trait, *f_w_* is the frequency of remaining ancestral phenotypes in the population). This shift from a time-slice view to a trait-centered view can lead to larger qualitative differences in particular if the mutation rate is low (Θ_*l*_ ≪ 1/*L*). In this case, a distribution of genetic architectures at a fixed time *t* > 0 will incorporate opposite cases where adaptation of the trait has either already been completed or not even started because the population still waits for a successful mutant. Biologically, a trait-centered view seems to be closer to the idea of an “architecture of phenotypic adaptation”. Mathematically, the changed perspective enables the derivation of analytical results. By comparing architectures for variable degrees of phenotypic adaptation, we still obtain a view of the adaptation dynamics, with *f_w_* as dynamical variable instead of time *t*. This is shown in Fig S.9. For Θ_*bg*_ ≤ 1, we see how the dominant contribution of a single “major locus” to the adaptive response emerges early on and then accentuates during the adaptive phase.

**Fig S.9.**
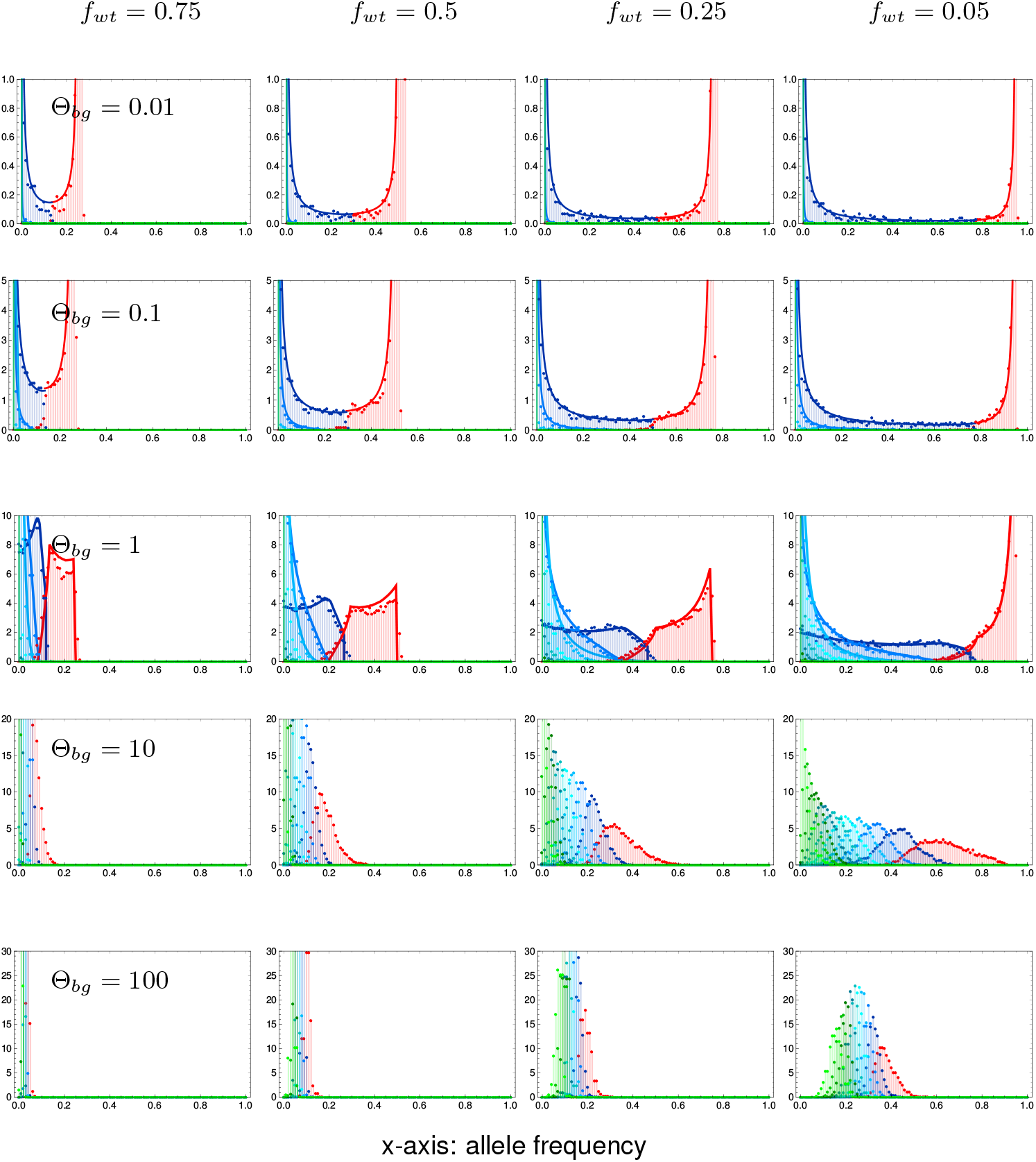
Dynamics of the adaptive process. Allele frequency distributions at four stages over the course of adaptation. Approximations correspond to the Fig S.7 each rescaled to the changed stopping condition *f_wt_* = 0.75; 0.5; 0.25; 0.05. Simulations for 10 000 replicates per mutation rate with *s_b_* = –*s_d_* = 0.1.

## Acknowledgments

We thank Claus Vogl for his insightful comments and several fruitful discussions. We also thank Matthias Maschek for his help concerning programming and simulation setup Finally, a special thank you goes to Montgomery Slatkin for his hospitality in welcoming JH and IH to his lab at UC Berkeley, where this project was started. We also thank Justin C. Fay and three anonymous referees for helpful comments and suggestions.

## Author Contributions

Authors: Ilse Höllinger (IH), PleuniS Pennings (PSP), Joachim Hermisson (JH)

1. JH, PSP and IH designed the study concept.
2. IH wrote the first version of the manuscript.
3. IH prepared the figures.
4. JH revised the manuscript with comments by PSP and IH.
5. JH derived the analytical results with input from PSP and IH.
6. IH wrote the simulations code with input from JH and PSP.
7. IH prepared the numerical results.

## Data Archiving

We provide a comprehensive *Mathematica* [63] notebook, showing visualizations of the analytical predictions. The simulation code and data, and summary statistics underlying all figures are available via Dryad.

Höllinger I, Pennings PS, Hermisson J. Data from: Polygenic adaptation: From sweeps to subtle frequency shifts. Dryad Digital Repository.

DOI: https://doi.org/10.5061/dryad.7n6vg10

## B Supplement

**Fig S.10.**
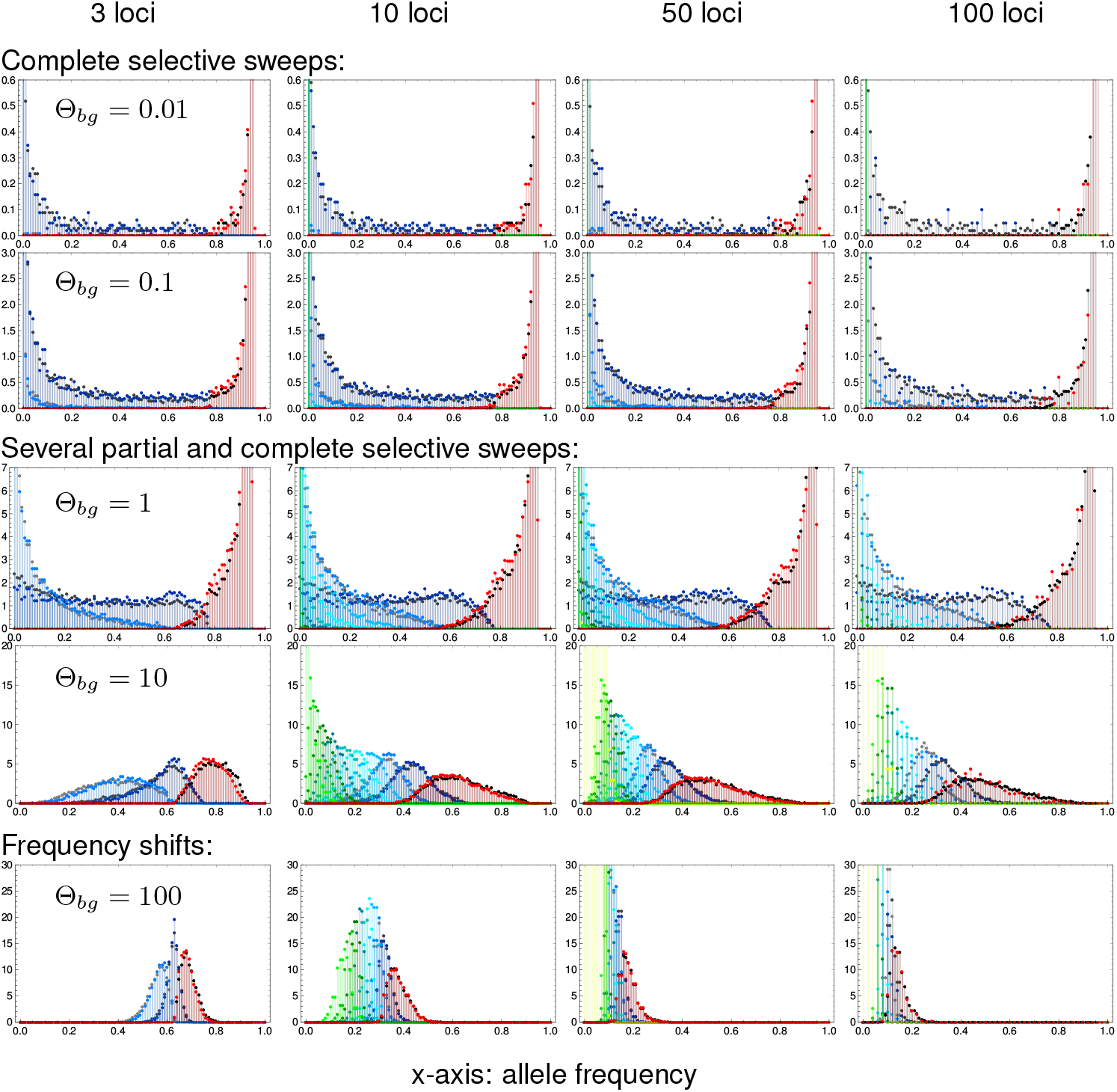
Weakly relaxed redundancy. Weakly relaxing redundancy such that a single mutant has fitness 1 + 0.9*s_b/d_* and only two mutations or more confer the full fitness effect (1 + *s_b/d_*) demonstrates the robustness of our model. As in Fig 4, allele frequency distributions of derived alleles are displayed once the frequency of the wildtype individuals in the population has decreased to *f_w_* = 5%, which corresponds to an increase of 95% in mean fitness for complete redundancy. Genomic patterns of adaptation show very similar characteristics as with complete redundancy. Simulation data for relaxed redundancy (colored dots) are almost identical to results for complete redundancy (gray dots).

**Fig S.11.**
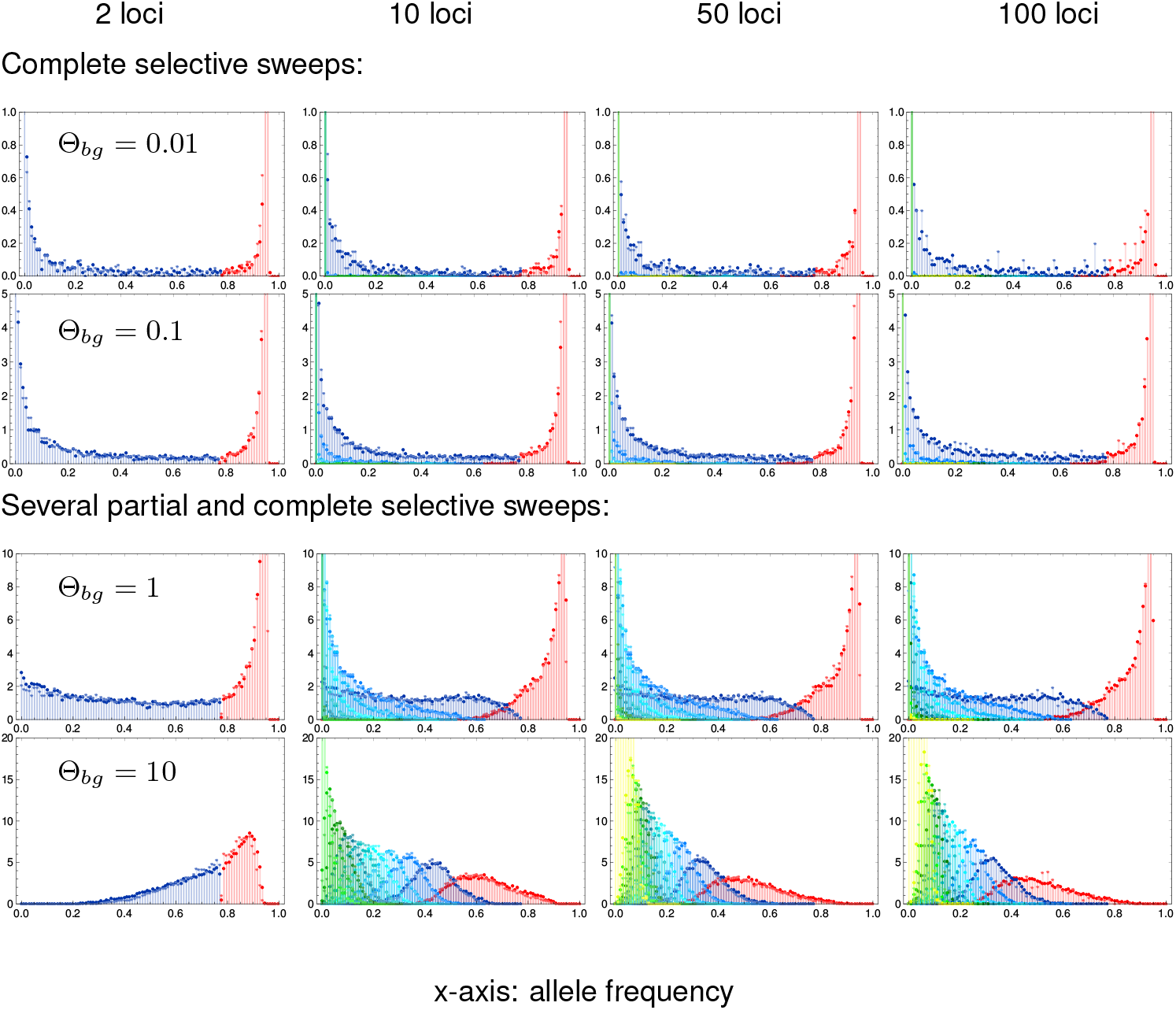
Genetic architecture with weak selection. Frequency distributions of major and minor loci are shown upon an increase of 95% in mean fitness for complete redundancy for *s_b_* = 0.1 (colored dots, data as in Fig 4) and weaker selection *s_b_* = 0.0] (colored asterisks). Deleterious selection before the environmental change is set to *s_d_* = –*s_b_*. As we condition on adaptation from the ancestral state, we do not obtain enough valid runs for *s_d_* = –0.01 and θ_*bg*_ = 100.

### Mathematical Appendix

This Appendix describes the details of the mathematical model and methods used to derive the analytical results of the article. Section M.1 gives an outline of the model; section M.2 introduces the branching process method used for the early stochastic phase of polygenic adaptation; section M.3 describes the derivation of the joint frequency distribution at the end of the deterministic phase.

#### M.1 Redundant trait model

Consider a panmictic population of *N_e_* haploids. Selection acts on a binary trait *Z* (e.g. resistance) with just two states, a wildtype state *Z*_0_ (not resistant) and a mutant state *Z*_1_ (resistant). Without restriction, we can choose *Z*_0_ = 0 and *Z*_1_ = 1. Malthusian (logarithmic) fitness is defined by the function

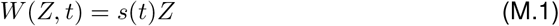

where the time dependent coefficient *s*(*t*) defines the strength of directional selection. We assume that *s*(*t*) < 0 for *t* < 0, but *s*(*t*) > 0 for *t* > 0, such that the optimal trait value shifts from the wildtype state *Z* = 0 to the mutant state *Z* = 1 due to some change in the environment at time *t* = 0. We also assume that selection is stronger than drift, |*Ns*(*t*)| ≫ 1 for almost all t, but is arbitrary otherwise.

We assume that *Z* is polygenic, with *L* biallelic loci (wildtype *a_i_* and mutant allele *A_i_, i* = 1,…, *L*) constituting its genetic basis. While genotype a = (*a*_1_, *a*_2_,…, *a_L_*) produces the ancestral wildtype *Z*_0_, all mutant genotypes are fully redundant and produce the mutant phenotype *Z*_1_, independently of the number of mutations. New mutations from *a_i_* to *A_i_* occur at a rate *μ_i_* per generation, with *μ_i_* ≪ |*s*(*t*)| for almost all *t*. For the purpose of our model, back mutation from *A_i_* to *a_i_* can be ignored. The linkage map among loci is arbitrary – unless explicitly specified otherwise. Let *p_i_* be the frequency of allele *A_i_*, and let *f_a_* be the frequency of the wildtype genotype a. Then the mean fitness in the population is

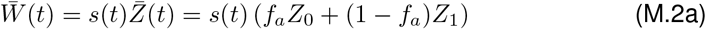

where 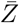 is the trait mean. Since *W*(*Z*_1_, *t*) = *s*(*t*)*Z*_1_ is the marginal fitness of any mutant allele, the selection dynamics at the ith locus can be expressed as

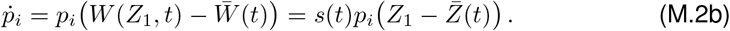

Our redundancy assumption implies strong diminishing returns epistasis on the level of fitness: the fitness of genotypes with multiple mutations is the same as the one of single mutants. Eq (M.2b) shows that the epistatic effect of the genetic background on the dynamics at a particular locus is mediated by the trait mean 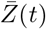 as single compound parameter. Allele frequencies at all loci change with the same (time and frequency-dependent) rate. We readily establish that

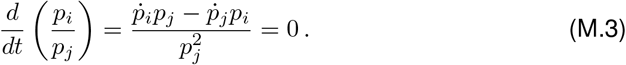

Thus, the ratio of allele frequencies among loci does not change under selection. Note that this holds for an arbitrary linkage map. We can conclude that any differences in (relative) allele frequencies are due to mutation and drift.

We are interested in the pattern of allele frequency changes across loci during the phase of rapid phenotypic adaptation. This phase starts with the onset of positive selection on derived alleles at time *t* = 0. It ends when mean fitness 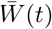 approaches its maximum *s*(*t*)*Z*_1_ and further selective change in the allele frequencies is strongly decelerated. Since 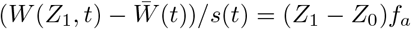, we can parametrize this end point by a condition *f_a_*(*t*) = *f_w_* on the frequency of the wildtype *Z*_0_ in the population. In our figures, we usually use *f_w_* = 0.05. As initial state at time *t* = 0, we assume that the population adapts from a balance of mutation, selection, and drift. We thus allow for standing genetic variation (SGV) at all loci. If selection prior to *t* = 0 is constant (which is what we generally assume in our computer simulations, see main text), SGV is given by the standard equilibrium distribution under mutation, selection, and drift, where we require that *a_i_* is the ancestral state at each locus. I.e., each allele frequency trajectory *p_i_*(*t*), back in time, originates from the boundary *p_i_* = 0 rather than *p_i_* = 1 (see also [1] for this concept). However, our analytical results do not require a static equilibrium and, for a general *s*(*t*) < 0 for *t* < 0, the SGV reflects this non-equilibrium dynamics.

As described in the main text, we dissect the adaptive process into two phases. During an initial *stochastic phase* mutation, selection, and drift lead to the build-up of genetic variation, either from SGV or due to new mutation after time *t* = 0, as long as allele frequencies *p_i_* at all loci are still low. We will describe our approach to this phase in detail in the section on Yule processes below. Once allele frequencies are sufficiently large, genetic drift and recurrent new mutation play only a minor role relative to selection until we reach the end of the rapid adaptive phase. We thus enter a *deterministic phase* where the dynamics is then well approximated by Eq (M.2b).

##### Relaxed redundancy

To relax the stringent redundancy condition of our model, it is natural to assume that a single mutation is not sufficient to produce the full mutant phenotype *Z*_1_ = 1, but only a partial phenotype *Z_q_* = *q* with 0 < *q* < 1. This makes the marginal fitness of mutant alleles dependent on the genetic background. If genotypes with two or more mutations produce *Z*_1_, we have

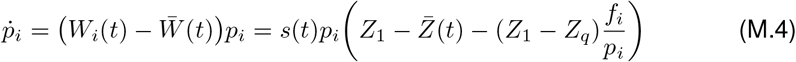

where *f_i_* is the frequency of the haplotype with a single mutation at locus *i*. Since *f_i_*/*p_i_* depends on *i* (even in linkage equilibrium), the ratio of allele frequencies at different loci is no longer invariant and the key symmetry assumption (M.3) of the fully redundant model is violated. Note that redundancy is recovered for very low mutant frequencies, such that double mutants are rare (*f_i_* ≈ *p_i_*) and also late in the adaptation process, when most haplotypes carry at least one mutation and *f_i_* → 0.

##### Diploids

We can generalize the redundant trait model to diploids as follows. For a general model, the dynamical equations in continuous time read

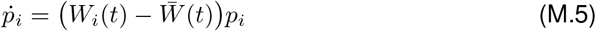

where *W_i_*(*t*) is the marginal fitness of allele *A_i_* and 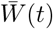 the mean fitness. All fitnesses may depend on the allele frequencies and on time. Using (M.3), we see that all mutant alleles *A_i_* are redundant in the sense that they all feel the same selection pressure if and only if their marginal fitnesses are equal at all times, *W_i_*(*t*) = *W_j_*(*t*), ∀ *i,j*. (The same condition can also be derived from a discrete time dynamics.) For haploids, equal marginal fitnesses, independently of the genetic composition of the population, enforces the fully redundant trait model described above. For diploids with dominance, the marginal fitness also depends on the allele frequency at the focal locus itself. An obvious solution to the condition of equal marginal fitnesses across loci is the case of complete dominance of the mutant allele. We can gain some more flexibility for the fitness scheme, if we assume that genotype frequencies are at Hardy-Weinberg equilibrium at all times. We can then distinguish three genotype classes: the wildtype without any mutations (normalized fitness 0), mutant individuals with one or more mutations on only a single haplotype (fitness *s*_1_(*t*)) and individuals with mutations on both haplotypes (fitness *s*_2_(*t*)). The marginal fitness of any mutant allele then is

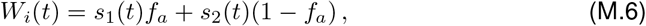

where *f_a_* is the frequency of the ancestral haplotype without mutations. We thus require redundancy of mutations (only) within haplotypes. Note, however, that this fitness scheme implies a position effect, i.e., the fitness of the genotype does not only depend on the number of mutations at each locus, but also on the association of mutations to one or the other haplotype. If we assume linkage equilibrium in addition to Hardy-Weinberg proportions, a position effect can be avoided if we use the following fitness scheme

1. The ancestral genotype without any mutants has normalized fitness *W*(*t*) = 0,
2. any genotype with at least one homozygous mutant has fitness *W*(*t*) = *s*_2_(*t*),
3. a genotype without a locus that is homozygous for the mutant, but with *k* loci that are heterozygous has fitness

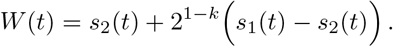 Since 2^1−*k*^ is the probability for any focal mutant allele to be on the same haplotype with all *k* – 1 other mutant alleles, assuming linkage equilibrium, this fitness scheme leads to the same marginal fitness as Eq (M.6) above.

#### M.2 Yule approximation

We describe the dynamics of mutant types at the different loci during the stochastic phase by a *multi-type Yule pure birth process with immigration*. Our framework builds on established mathematical theory [2,3] and a previous approach to describe the genealogy of a beneficial allele during a selective sweep in terms of a Yule process [4,5]. Here, we extend this approach to the polygenic scenario.

Consider a mutation *A_i_* that appears at some locus either prior to the environmental change (standing genetic variation) or after the change. This mutation is relevant for the joint distribution of mutant allele frequencies at the time of observation after the rapid adaptive phase if and only if descendants of this mutation still segregate in the population at this time. The idea of the Yule approach is to construct the genealogies of these mutant descendants at all loci forward in time. We start the process at some time *t*_0_ ≪ 0 in the past before the first mutation with surviving descendants has originated. We assume that the frequency *p_i_* of mutant alleles is low during the entire stochastic phase. Then, new mutations at locus *i* appear at rate ≈ *N_μ_i__* =: Θ_*i*_/2 per generation, but only a fraction of those will survive deleterious selection prior to *t* = 0 and genetic drift to establish in the population and to contribute to the adaptation of the trait. We denote this establishment probability as *p*_est_(*t*). If selection is constant and positive (as assumed in the main text), *s*(*t*) = *s_b_* > 0, we can approximate *p*_est_ ≈ 2*s_b_*. For general time-dependent selection, *p*_est_(*t*) will depend on 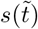 with 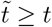 [6], and also on the mutations that were previously established at the same or at other loci. Crucially, however, since the marginal fitness of mutant copies at all loci is the same at any given time, *p*_est_(*t*) does not depend on the locus. We only include mutants into our Yule process that successfully establish in the population, which are represented as “immortal lineages” in the Yule tree. We follow these lineages in continuous time. There are then two types of events:

1. First, new mutation creates new immortal lineages at rate

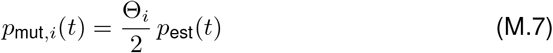

independently at each locus. This event is called “immigration” in the mathematical literature [2], but it corresponds to mutation in our model. (In a model with gene flow, where adaptation in a local deme occurs from immigration, new lines would be truly immigrants, see also [7] for this analogy).
2. Second, existing immortal mutant alleles *A_i_* can give birth to further immortal mutant copies, corresponding to a split of the immortal line in the Yule process. To derive the split rate *p*_split_, imagine that we implement the evolutionary dynamics as a continuous-time Moran model, where individuals give birth (due to a binary split) at constant rate one per generation. In the corresponding Yule process, we only include this birth event if it leads to two immortal lineages. Obviously, the probability to “be immortal” for a newborn individual is the same as for a new mutation and given by *p*_est_(*t*). Conditioning on the fact that we only consider splits of immortal lineages and thus at least one of the offspring lineages must be immortal, we arrive at a split rate per immortal lineage of

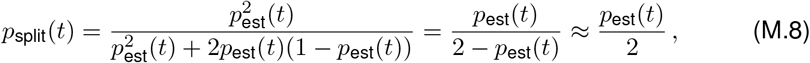

where the approximation in the last term assumes that *p*_est_(*t*) ≪ 1, which is usually the case unless selection is very strong.

The Yule process defines a continuous-time Markov process of a random variable k = (*k*_1_,…, *k_L_*), where 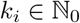 is the number of immortal mutant lineages at the *i*th locus. We are interested in the relative proportions in the number of lineages *k_i_* across loci after a sufficiently long time – assuming that the distribution of these proportions reaches a limit by the end of the stochastic phase. We can generate this distribution from the transition probabilities among Yule states (the embedded jump-chain of the continuous-time process). If there are currently (*k*_1_,…, *k_L_*) lineages at the *L* loci, the probability that the next event is either a birth event (split) or a new mutation (immigration) at locus *i* is

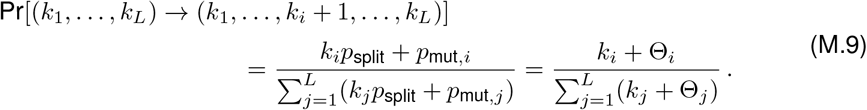

Crucially, these transition probabilities are constant in time and independent of the establishment probability *p*_est_(*t*). As a consequence, they are also independent of the mutant fitness, which only affects the speed of the Yule process (via *p*_est_), but not its sequence of events.

We start the process with no mutants and stop it whenever the number of mutants at one of the loci(e.g. locus 1) reaches some number *k*_1_ = *n*. We are interested in the distribution of the number of mutants *k_i_* at the other loci at this time, respectively their ratios *k_i_*/*n* (remember that we already know that these ratios stay invariant during the deterministic phase of the adaptation process). We can prove the following

##### Theorem 1

In the limit of *n* → ∞, the joint distribution of ratios *x_i_* = *k_i_*/*n* of immortal mutant lineages across loci converges to the *inverted Dirichlet distribution*,

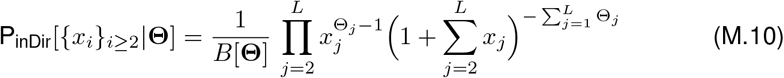

where the vector **Θ** = (Θ_1_,…, Θ_*L*_) summarizes the mutation rates and *B*[**Θ**] is the multivariate Beta function, which can be expressed in terms of Gamma functions as

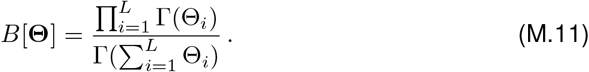

##### Proof

We proceed in three steps.

**Step 1** Assume that we stop the process when the first locus reaches *n* > 0 lineages. We derive the probability that the process at this time is in state (*n, k*_2_,…, *k_L_*) as follows. We need *n* + *k*_2_ + … + *k_L_* events (new mutations or splits) to generate all mutant individuals. The last event must occur at the first locus. All other events can occur in arbitrary order at the *L* loci. The probability of each realization (each order of events at the loci) is given by the corresponding product of transition probabilities (M.9) The key insight is that all realizations have the same probability. Indeed, the denominator of (M.9) does not depend on the locus where the next event occurs. Different realizations then only correspond to permutations in the factors *k_i_* + Θ_*i*_ in the numerator of the product of transition probabilities. We can directly write down the probability for the state as

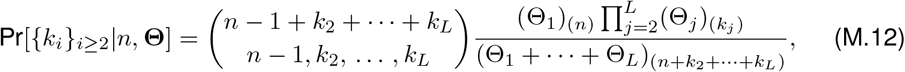

where

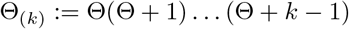

is the Pochhammer function. The leading multinomial coefficient counts the number of all permutations and the ratio of Pochhammer functions is the probability of each realization.

**Step 2** We can rewrite (M.12) as a *Dirichlet-negative-multinomial* compound distribution, defined as

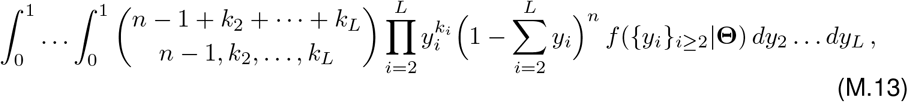

where

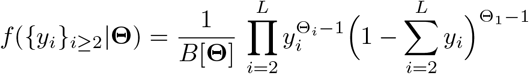

is the (*L* – 1)-dimensional Dirichlet distribution for a *L*-dimensional probability vector (*y*_1_,…, *y_L_*) with constraint 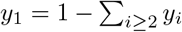. This is best shown in the reverse direction, i.e., by deriving (M.12) from (M.13). To see this, note that

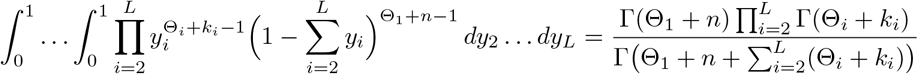

because the integrand in this expression is just a Dirichlet density with shifted values of Θ_*i*_ → Θ_*i*_ + *k_i_* and the right hand side is the corresponding normalization factor. Then using

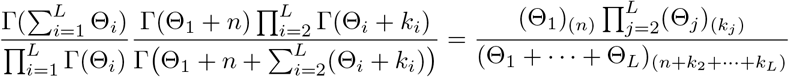

reduces (M.13) to (M.12).

The compound distribution Eq (M.13) can be interpreted as follows: If a random experiment can have a finite number of outcomes (here: mutant lineages at one of *L* loci), the negative multinomial distribution describes the probability to observe each of these events *k_i_* times if we repeat the experiment until a focal event (here: new mutant lineage at the first locus) has occurred *n* times. While the negative multinomial distribution assumes that all outcomes occur with a fixed probability *y_i_*, this probability is itself drawn from a Dirichlet distribution in the Dirichlet-negative-multinomial compound distribution. In the present context, the main advantage of (M.13) over (M.12) is that we can easily perform the limit *n* → ∞ in this form.

**Step 3** For large *n* → ∞, the values of *k_i_*/*n, i* ≥ 2, of the negative multinomial distribution can be replaced by their expectations,

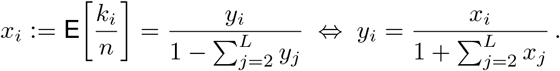

We can then transform the density (M.10) from variables *y_i_* to the *x_i_* (representing the relative mutant frequencies). The entries of the Jacobian matrix (for 2 ≤ *i,j* ≤ *L*) are

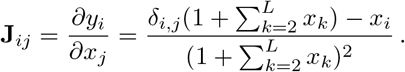

Since this is the sum of an identity matrix (times a factor) and a matrix with identical columns we can easily derive the eigenvalues and thus the determinant,

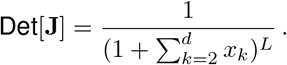

Applying this transformation to (M.13), we obtain (M.10).

##### Remarks

1. For two loci, the Dirichlet-negative-multinomial distribution (M.13) reduces to a *Beta-negative-binomial* distribution

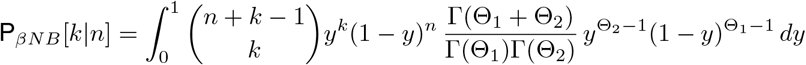

and the inverted Dirichlet distribution (M.10) simplifies to a so-called *β-prime* distribution,

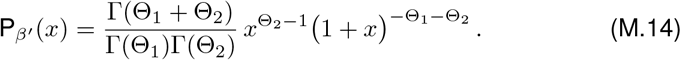 If we measure the ratio *x* always relative to the locus with the higher frequency, we obtain a conditioned distribution that is truncated at *x* = 1. For equal locus mutation rates Θ_1_ = Θ_2_ = Θ_*l*_, in particular,

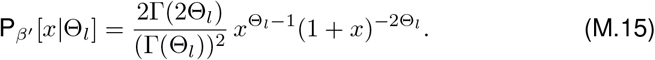

with expectation

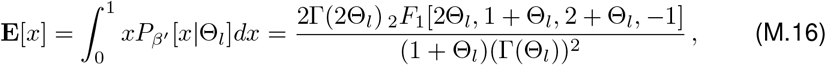

where _2_*F*_1_ is the hypergeometric function.
2. The process described here is a variant of the *Polya urn* and *Hoppe urn* processes that are well-known in the mathematical literature and have been used to describe coalescent processes forward in time [2,3].
3. Our result (M.10) can also be seen as multi-locus version of Wright’s formula for the stationary distribution of the Wright-Fisher diffusion [8]. For *L* neutral alleles at a singe locus, and if the mutation rates θ_*i*_ depend only on the target allele (house-of-cards condition), this is a Dirichlet distribution. Here, we see that an analogous result holds for a distribution of equivalent (mutually redundant) alleles across *L* loci. Although alleles at different loci cannot mutate into each other and are never identical by descent, it turns out that the genealogy in both models can be described by a Yule process with immigration. In contrast to the single-locus case, we obtain an *inverted* Dirichlet distribution for multiple loci. This difference results from a different stopping condition for the Yule process. For a single locus, the population size sets an upper bound for the total number of copies across all alleles. If we stop the process for a given total number *n*_tot_ of lines, we obtain the classical Dirichlet distribution in the limit *n*_tot_ → ∞. In contrast, the population size defines a bound for mutants of a only single type in the multi-locus case, which is reflected by our choice of the stopping condition. This choice is appropriate unless all loci are tightly linked, as we will see below.
4. In our model, we did not distinguish different mutational origins of mutant alleles at the same locus. It is, in principle, possible to do so. For any single locus, the process *conditioned on* reaching some number of mutants *k_i_* at this locus *i* is entirely independent of the process at the other loci. The joint distribution of different mutational origins at this locus is therefore given by the Ewens sampling formula, as described in the theory of soft selective sweeps ([7,9]).

#### M.3 Allele frequency distributions

Eq (M.10) predicts the distribution of allele frequency ratios *x_i_* at the end of the stochastic phase of the adaptive process. Typically, the Yule process will approach convergence for *n* ≿ 100. In a large population, this still corresponds to a small allele frequency. However, since the allele frequency ratios remain constant also during the deterministic phase, we can use the Yule process result to derive the distribution of mutant allele frequencies also at a later stage, when (partial or complete) phenotypic adaptation has been achieved. As above, we characterize the time of observation via the frequency of the ancestral phenotypes *f_w_* that is still found in the population. We treat the case of full adaptation, *f_w_* = 0, before we turn to the case of a general *f_w_*.

##### Complete phenotypic adaptation, *f_w_* = 0

If selection is very strong, complete fixation of the mutant phenotype may be rapidly achieved. For any non-zero level of recombination among loci, *f_w_* = 0 requires, in our model, that there is (at least) a single locus where the mutant allele has reached fixation. In the following, we will call the locus with the largest mutant frequency the *major locus* and all other loci *minor loci*. We are interested in the joint distribution of allele frequencies when the major locus has reached fixation. From (M.10), we can derive the probability that the first locus ends up being the major locus as

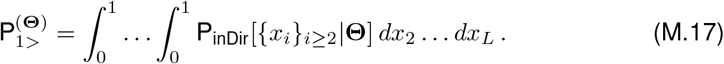

Since allele frequencies *p_i_* equal allele frequency ratios *x_i_* relative to the major locus in this case, the joint distribution at all minor loci, {*p_i_*}_*i*≥2_, 0 ≤ *p_i_* ≤ 1, conditioned on fixation of the mutant allele at the first locus, follows as P_inDir_[{*p_i_*}_*i*≥2_|Θ]/P_1>_[Θ]. The joint allele frequency distribution for all loci at *f_w_* =0 results as product of a Dirac point measure at the major locus and truncated inverted Dirichlet densities at the minor loci. Summing over all possible loci as major locus we obtain

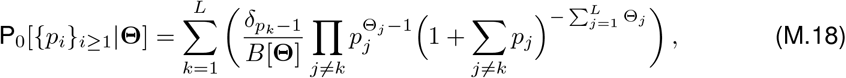

where the Dirac *δ* constrains the distribution to the boundary faces *p_k_* = 1 of the *L*-dimensional hypercube [0,1]^*L*^ of allele frequencies.

Note that this formula is independent of linkage patterns as long as loci can recombine at all and are not completely linked (see below for this case).

##### Incomplete phenotypic adaptation, *f_w_* > 0, linkage equilibrium

While the distribution of allele frequency *ratios x_i_*, Eq (M.10), holds for any time of observation during the adaptive process (once the Yule process has reached convergence), the corresponding distribution (M.18) for the *absolute* allele frequencies *p_i_* holds only for complete phenotypic adaptation, *f_w_* = 0. To derive this distribution for arbitrary *f_w_* ≥ 0, we need to translate the stopping condition for the ancestral phenotype to a condition on the *p_i_*. For *f_w_* = 0, this just leads to the condition *p_k_* = 1 for the major locus, constraining the distribution (M.18) to the boundary faces of the allele frequency hypercube. Importantly, this constraint is independent of linkage. For *f_w_* > 0, in contrast, any constraint on the distribution of the *p_i_* due to the stopping condition will necessarily also depend on the linkage disequilibria. For further analytical progress we now assume that recombination is sufficiently strong that linkage disequilibria can be ignored. We then obtain

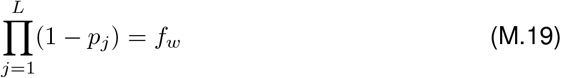

and the joint allele frequency distribution is given by the following Theorem, which is our main analytical result.

###### Theorem 2

If the adaptive process is stopped at a frequency *f_w_* of the ancestral phenotype in the population, and assuming linkage equilibrium among loci, the joint distribution of mutant frequencies on the *L*-dimensional hypercube is

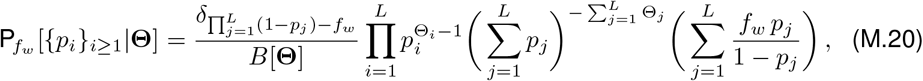

where the *δ*-function restricts the support of P_*fw*_ [{*p_j_*}_*i*≥1_|Θ] to the (*L* – 1)-dimensiona submanifold 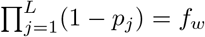.

###### Proof

We can rewrite (M.19) as condition on the frequency *p*_1_ at the first locus,

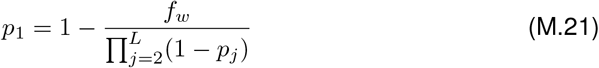

to obtain the transformation from frequency ratios *x_i_* to absolute allele frequencies *p_i_*, *i* ≥ 2,

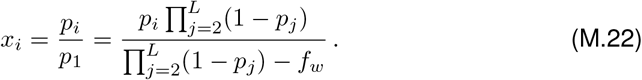

The corresponding Jacobian matrix reads (2 ≤ *i,j* ≤ *L*)

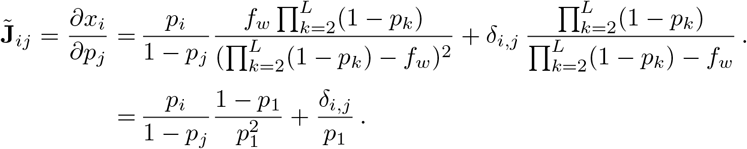

Thus

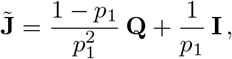

where **I** is the identity matrix and **Q**_*i,j*_ = *p_i_*/(1 – *p_j_*). Since **Q** has the eigenvalue 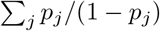 and a (*L* – 2)-fold eigenvalue 0, we obtain the spectrum of 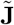 and thus the determinant

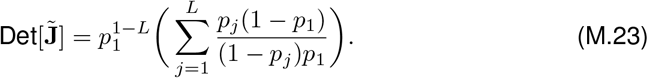

From (M.10), we then obtain the joint distribution of locus frequencies *p*_2_,…, *p_L_* at the stopping condition (M.21) as

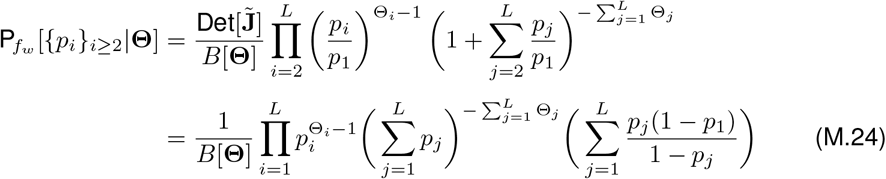

where the dependence on *f_w_* is implicit in *p*_1_ = *p*_1_(*f_w_*), as given in (M.21). The joint distribution over all *L* locifollows as

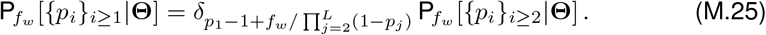

Note that we do not assume that the first locus is the major locus in (M.25). Finally, the symmetrical form (M.20) results from the relation

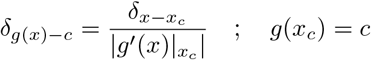

for the Dirac *δ*-function.

###### Remarks

1. To obtain marginal distributions for single lociwe generally need to perform a (*L* – 2)-dimensional integral (after resolving the *δ*-function). Details for specific cases used in the main part of the article are provided in the Mathematica notebook. For two loci, simple explicit formulas for marginal distributions can be derived. E.g., the marginal distribution at the first locus reads

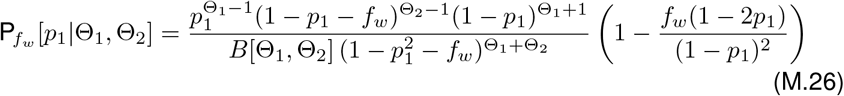

for 0 ≤ *p*_1_ ≤ *f_w_*. The distribution has singularities at *p*_1_ = 0 for Θ_1_ < 1 and at *p*_1_ = 1 – *f_w_* for Θ_2_ < 1. The distributions 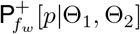 at the major locus and 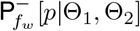 at the minor locus (which can either be locus 1 or locus 2) follow as

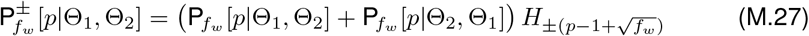

where *H*(*x*) is the Heaviside function with *H_x_* = 1 for *x* ≥ 0 and *H_x_* = 0 else. Finally, the *conditioned* distributions 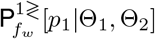 at the first locus if this locus is the major/minor locus are

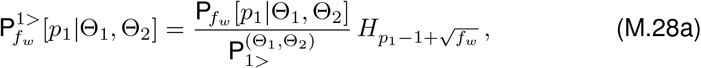

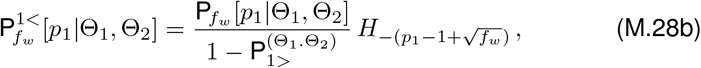

where 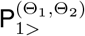, defined in Eq (M.17), evaluates to a Hypergeometric function for general Θ_1_ ≠ Θ_2_, but reduces to 1/2 for Θ_1_ = Θ_2_.
2. The marginal distribution for *p_k_* has a singularity at *p_k_* = 0 for *Θ_k_* < 1 and a singularity at *p_k_* = 1 – *f_w_* for 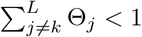. To see this, consider the marginal distribution of *p_L_*, which is obtained from Eq. (M.25) after integartion over *p*_1_,…, *p*_*L*–1_. Dropping non-singular terms (such as the sums in Eq M.24), and defining

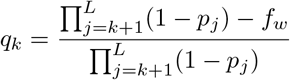

the singlular part can be written as

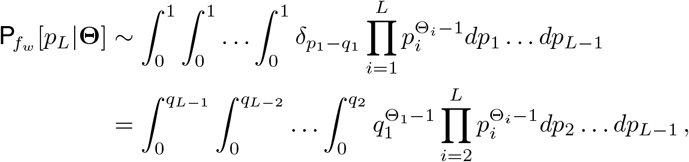

after performing the *p*_1_ integral. The upper integral limits *q_k_* account for the constraint *q*_1_ > 0. Substituting

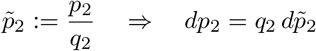

and using that 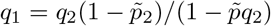 we obtain

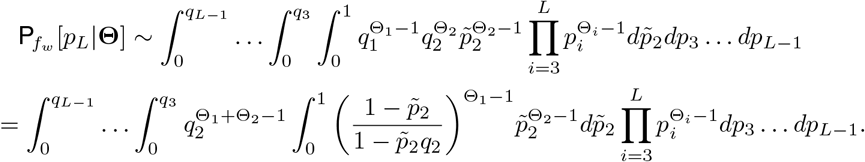 Since the 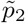 integral is bounded by 1/Θ_2_ from below and by 1/Θ_2_ + 1/Θ_1_ from above for all 0 ≤ *q*_2_ ≤ 1, it does not contribute to a singularity in *P_f_w__*[*p_L_*|Θ]. For the singular part, we thus have

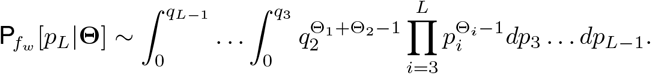 Iterating the substitution procedure for variables *p*_3_ to *p*_*L*–1_, we arrive at

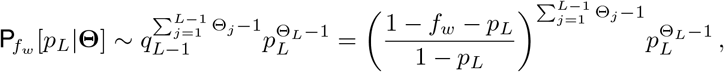

demonstrating the singular behavior for *p_L_* → 0 and for *p_L_* → 1 – *f_w_*. Since the labeling of loci is arbitrary, the assertion follows for all loci.

##### Incomplete phenotypic adaptation, *f_w_* > 0, tight linkage

Even if all loci are completely linked, the joint distribution of allele frequency *ratios* is still given by (M.10). However, the transformation to absolute allele frequencies at the stopping condition *f_w_* ≠ 0 depends on linkage. Because all mutant alleles are rare during the stochastic phase, we can ignore haplotypes with more than a single mutant during this time. Since we ignore new mutations during the deterministic phase, mutant alleles stay in maximal linkage disequilibrium in the absence of recombination. We thus have

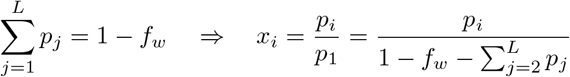

with corresponding Jacobian

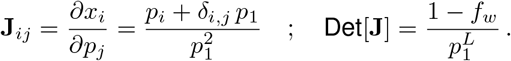

Using this transformation on (M.10), the joint distribution of mutant frequencies reads

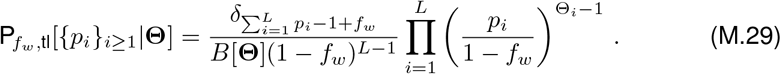

Evidently, this is just the Dirichlet distribution on the cube [0,1 – *f_w_*]^*L*^. This is expected since the problem reduces to a single-locus, *L*-alleles problem for tight linkage. The marginal distributions can be derived for an arbitrary number of loci and are given by transformed *β*-distributions,

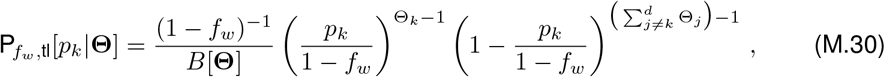

with singularities at the boundaries *p_k_* = 0 for Θ_*k*_ < 1 and at *p_k_* = 1 – *f_w_* for 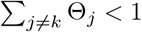 as in the linkage equilibrium case. For two tightly linked loci, the major locus must have frequency *p* > (1 – *f_w_*)/2. The distribution at the major/minor locus therefore reads

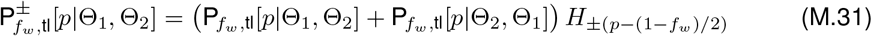

and conditioned distributions follow as in (M.28).

## References

1. Barton NH, Keightley PD. Multifactorial genetics: understanding quantitative genetic variation. Nature Reviews Genetics. 2002;3(1):11–21.

2. Messer PW, Ellner SP, Hairston Jr NG. Can population genetics adapt to rapid evolution? Trends in Genetics. 2016;32(7):408–418.

3. Maynard-Smith J, Haigh J. The hitch-hiking effect of a favourable gene. Genetics Research. 1974;23(1):23–35.

4. Kaplan NL, Hudson R, Langley C. The ‘‘hitchhiking effect” revisited. Genetics. 1989;123(4):887–899.

5. Barton NH. The effect of hitch-hiking on neutral genealogies. Genetics Research. 1998;72(2):123–133.

6. Hermisson J, Pennings PS. Soft sweeps and beyond: understanding the patterns and probabilities of selection footprints under rapid adaptation. Methods in Ecology and Evolution. 2017;8(6):700–716.

7. Pritchard JK, Di Rienzo A. Adaptation-not by sweeps alone. Nature Reviews Genetics. 2010;11(10):665–667.

8. Boyle EA, Li YI, Pritchard JK. An expanded view of complex traits: from polygenic to omnigenic. Cell. 2017;169(7):1177–1186.

9. Fisher R. The correlation between relatives on the supposition of Mendelian Inheritance. Trans Roy Soc Edinburgh. 1918;52:339–433.

10. Barton NH, Etheridge A, Veber A. The infinitesimal model: Definition, derivation, and implications. Theoretical Population Biology. 2017;118:50–73.

11. Pritchard JK, Pickrell JK, Coop G. The genetics of human adaptation: hard sweeps, soft sweeps, and polygenic adaptation. Current Biology. 2010;20(4):208–215.

12. Hancock AM, Alkorta-Aranburu G, Witonsky DB, Di Rienzo A. Adaptations to new environments in humans: the role of subtle allele frequency shifts. Philosophical Transactions of the Royal Society of London B: Biological Sciences. 2010;365(1552):2459–2468.

13. Berg JJ, Coop G. A population genetic signal of polygenic adaptation. PLoS Genetics. 2014;10(8):e1004412.

14. Field Y, Boyle EA, Telis N, Gao Z, Gaulton KJ, Golan D, et al. Detection of human adaptation during the past 2000 years. Science. 2016;354(6313):760–764.

15. Csillery K, Rodríguez-Verdugo A, Rellstab C, Guillaume F. Detecting the genomic signal of polygenic adaptation and the role of epistasis in evolution. Molecular Ecology. 2018;27(3):606–612.

16. Berg JJ, Harpak A, Sinnott-Armstrong N, Joergensen AM, Mostafavi H, Field Y, et al. Reduced signal for polygenic adaptation of height in UK Biobank. bioRxiv. 2018;doi:10.1101/354951.

17. Sohail M, Maier RM, Ganna A, Bloemendal A, Martin AR, Turchin MC, et al. Signals of polygenic adaptation on height have been overestimated due to uncorrected population structure in genome-wide association studies. bioRxiv. 2018;doi:10.1101/355057.

18. Stephan W. Signatures of positive selection: from selective sweeps at individual loci to subtle allele frequency changes in polygenic adaptation. Molecular Ecology. 2016;25(1):79–88.

19. Turelli M, Barton NH. Dynamics of polygenic characters under selection. Theoretical Population Biology. 1990;38(1):1–57.

20. Turelli M, Barton NH. Genetic and statistical analyses of strong selection on polygenic traits: what, me normal? Genetics. 1994;138(3):913–941.

21. Burger R, Lynch M. Evolution and extinction in a changing environment: a quantitative-genetic analysis. Evolution. 1995;49(1):151–163.

22. Burger R. The mathematical theory of selection, recombination, and mutation. Wiley, Chichester, UK; 2000.

23. Geritz SA, Mesze G, Metz JA, et al. Evolutionarily singular strategies and the adaptive growth and branching of the evolutionary tree. Evolutionary Ecology. 1998;12(1):35–57.

24. Orr HA. The genetic theory of adaptation: a brief history. Nature Reviews Genetics. 2005;6(2):119–127.

25. Matuszewski S, Hermisson J, Kopp M. Catch me if you can: adaptation from standing genetic variation to a moving phenotypic optimum. Genetics. 2015;200(4):1255–1274.

26. Chevin LM, Hospital F. Selective sweep at a quantitative trait locus in the presence of background genetic variation. Genetics. 2008;180:1645–1660.

27. Lande R. The response to selection on major and minor mutations affecting a metrical trait. Heredity. 1983;50(1):47–65.

28. Pavlidis P, Metzler D, Stephan W. Selective sweeps in multilocus models of quantitative traits. Genetics. 2012;192(1):225–239.

29. Wollstein A, Stephan W. Adaptive fixation in two-locus models of stabilizing selection and genetic drift. Genetics. 2014;198(2):685–697.

30. Jain K, Stephan W. Response of polygenic traits under stabilizing selection and mutation when loci have unequal effects. G3: Genes, Genomes, Genetics. 2015;5(6):1065–1074.

31. Jain K, Stephan W. Rapid adaptation of a polygenic trait after a sudden environmental shift. Genetics. 2017;206(1):389–406.

32. Wright S. Evolution in Mendelian populations. Genetics. 1931;16(2):97–159.

33. Kryazhimskiy S, Rice DP, Jerison ER, Desai MM. Global epistasis makes adaptation predictable despite sequence-level stochasticity. Science. 2014;344(6191):1519–1522.

34. Ralph PL, Coop G. Parallel adaptation: one or many waves of advance of an advantageous allele? Genetics. 2010;186:647–668.

35. Ralph PL, Coop G. The role of standing variation in geographic convergent adaptation. The American Naturalist. 2015;186(S1):S5–S23.

36. Paulose J, Hermisson J, Hallatschek O. Spatial soft sweeps: patterns of adaptation in populations with long-range dispersal. (In Press) PLoS Genetics. 2019;.

37. Hermisson J, Pennings PS. Soft sweeps: molecular population genetics of adaptation from standing genetic variation. Genetics. 2005;169(4):2335–2352.

38. Etheridge A, Pfaffelhuber P, Wakolbinger A, et al. An approximate sampling formula under genetic hitchhiking. The Annals of Applied Probability. 2006;16(2):685–729.

39. Hermisson J, Pfaffelhuber P. The pattern of genetic hitchhiking under recurrent mutation. Electronic Journal of Probability. 2008;13:2069–2106.

40. Pennings PS, Hermisson J. Soft sweeps II-molecular population genetics of adaptation from recurrent mutation or migration. Molecular Biology and Evolution. 2006;23(5):1076–1084.

41. de Vladar HP, Barton NH. Stability and response of polygenic traits to stabilizing selection and mutation. Genetics. 2014;197(2):749–767.

42. Jain K, Devi A. Polygenic adaptation in changing environments. EPL (Europhysics Letters). 2018;123(4):48002.

43. Burger R, Gimelfarb A. Genetic variation maintained in multilocus models of additive quantitative traits under stabilizing selection. Genetics. 1999;152(2):807–820.

44. Coffman CJ, Doerge RW, Simonsen KL, Nichols KM, Duarte C, Wolfinger RD, et al. Model selection in binary trait locus mapping. Genetics. 2005;170:1281–1297.

45. Novembre J, Han E. Human population structure and the adaptive response to pathogen-induced selection pressures. Phil Trans R Soc B. 2012;367(1590):878–886.

46. Falconer D, Mackay T. Introduction to Quantitative Genetics. 4th ed. Longmans Green, Harlwo, Essex, UK; 1996.

47. Hill WG. Applications of population genetics to animal breeding, from Wright, Fisher and Lush to genomic prediction. Genetics. 2014;196(1):1–16.

48. Visscher PM, Wray NR, Zhang Q, Sklar P, McCarthy MI, Brown MA, et al. 10 years of GWAS discovery: biology, function, and translation. The American Journal of Human Genetics. 2017;101(1):5–22.

49. Laporte M, Pavey SA, Rougeux C, Pierron F, Lauzent M, Budzinski H, et al. RAD sequencing reveals within-generation polygenic selection in response to anthropogenic organic and metal contamination in North Atlantic Eels. Molecular Ecology. 2016;25(1):219–237.

50. Zan Y, Carlborg Ö. A multilocus association analysis method integrating phenotype and expression data reveals multiple novel associations to flowering time variation in wild-collected Arabidopsis thaliana. Molecular Ecology Resources. 2018; p. 798–808.

51. Karasov T, Messer PW, Petrov DA. Evidence that adaptation in *Drosophila* is not limited by mutation at single sites. PLoS Genetics. 2010;6(6):e1000924.

52. Barton NH. Understanding adaptation in large populations. PLoS Genetics. 2010;6(6):e1000987.

53. Wilson BA, Petrov DA, Messer PW. Soft selective sweeps in complex demographic scenarios. Genetics. 2014; p. 669–684.

54. Lazaridis I, Nadel D, Rollefson G, Merrett DC, Rohland N, Mallick S, et al. Genomic insights into the origin of farming in the ancient Near East. Nature. 2016;536(7617):419–424.

55. Pickrell JK, Reich D. Toward a new history and geography of human genes informed by ancient DNA. Trends in Genetics. 2014;30(9):377–389.

56. Yeaman S. Local adaptation by alleles of small effect. The American Naturalist. 2015;186(S1):S74–S89.

57. Franssen SU, Kofler R, Schlötterer C. Uncovering the genetic signature of quantitative trait evolution with replicated time series data. Heredity. 2017;118(1):42–51.

58. Burke MK, Dunham JP, Shahrestani P, Thornton KR, Rose MR, Long AD. Genome-wide analysis of a long-term evolution experiment with *Drosophila*. Nature. 2010;467(7315):587–590.

59. Barghi N, Tobler R, Nolte V, Jaksic AM, Mallard F, Otte K, et al. Genetic redundancy fuels polygenic adaptation in *Drosophila*. (In Press) PLoS Biology. 2019;.

60. Griffiths R, Tavaré S. The age of a mutation in a general coalescent tree. Stochastic Models. 1998;14(1-2):273–295.

61. Hoppe FM. Pólya-like urns and the Ewens’ sampling formula. Journal of Mathematical Biology. 1984;20(1):91–94.

62. Orr HA, Betancourt AJ. Haldane’s sieve and adaptation from the standing genetic variation. Genetics. 2001;157(2):875–884.

63. Wolfram Research I. Mathematica, Version 11.3;.

## References

1. Hermisson J, Pennings PS. Soft sweeps: molecular population genetics of adaptation from standing genetic variation. Genetics. 2005;169(4):2335–2352.

2. Joyce P, Tavare S. Cycles, permutations and the structure of the Yule process with immigration. Stochastic processes and their applications. 1987;25:309–314.

3. Durrett R. Probability: theory and examples. Cambridge University Press; 2010.

4. Etheridge A, Pfaffelhuber P Wakolbinger A, et al. An approximate sampling formula under genetic hitchhiking. The Annals of Applied Probability. 2006;16(2):685–729.

5. Hermisson J, Pfaffelhuber P. The pattern of genetic hitchhiking under recurrent mutation. Electronic Journal of Probability. 2008;13:2069–2106.

6. Uecker H, Hermisson J. On the fixation process of a beneficial mutation in a variable environment. Genetics. 2011;188(4):915–930.

7. Pennings PS, Hermisson J. Soft sweeps II-molecular population genetics of adaptation from recurrent mutation or migration. Molecular Biology and Evolution. 2006;23(5):1076–1084.

8. Wright S. Evolution in Mendelian populations. Genetics. 1931;16(2):97–159.

9. Hermisson J, Pennings PS. Soft sweeps and beyond: understanding the patterns and probabilities of selection footprints under rapid adaptation. Methods in Ecology and Evolution. 2017;8(6):700–716.

